# Experimental Diets Dictate the Metabolic Benefits of Probiotics in Obesity

**DOI:** 10.1101/2022.10.27.514016

**Authors:** Ida Søgaard Larsen, Béatrice S.-Y. Choi, Bandik Föh, Nanna Ny Kristensen, Adia Ouellette, Rune Falkenberg Haller, Peter Bjarke Olsen, Delphine Saulnier, Christian Sina, Benjamin A. H. Jensen, André Marette

**Author notes:** Co-senior authors (Benjamin A. H. Jensen, André Marette). Novozymes A/S, Denmark, Krogshoejvej 36, DK-2880 Bagsvaerd, Denmark. **Author Contributions** BAHJ, ISL, NNK, AM and BSYC conceived and designed the studies. ISL, BAHJ, and BSYC carried out the *in vivo* studies. ISL, BSYC, BF, AO, RFH, PBO, and DS generated data. BAHJ designed the diets and supervised all parts of the study; AM and CS supervised parts of the study. ISL integrated the data and wrote the first drafts of the manuscript, which was revised by all the authors, who agreed upon the submitted manuscript.

## Abstract

**Background:** Growing evidence supports the use of probiotics to prevent or mitigate obesity-related dysmetabolism and non-alcoholic fatty liver disease (NAFLD). However, frequent reports of responders versus non-responders to probiotic treatment warrant a better understanding of key modifiers of host-microbe interactions. The influence of host diet on probiotic efficacy, in particular against metabolic diseases, remains elusive.

**Method:** We fed C57BL6/J mice a low fat reference diet or one of two energy-matched high fat and high sucrose diets for 12 weeks; a classical high fat diet (HFD) and a customized fast food-mimicking diet (FFMD). During the studies, mice fed either obesogenic diet were gavaged daily with one of two probiotic lactic acid bacteria (LAB) strains previously classified as *Lactobaccillus*, namely *Limosilactobacillus reuteri* (*L. reuteri*) or *Lacticaseibacillus paracasei* subsp. *paracasei* (*L. paracasei*), or vehicle.

**Results:** The tested probiotics exhibited a reproducible efficacy but dichotomous response according to the obesogenic diets used. Indeed, *L. paracasei* prevented weight gain, improved insulin sensitivity, and protected against NAFLD development in mice fed HFD, but not FFMD. Conversely, *L. reuteri* improved glucoregulatory capacity, reduced NAFLD development, and increased distal gut bile acid levels associated with changes in predicted functions of the gut microbiota exclusively in the context of FFMD-feeding.

**Conclusion:** We found that the probiotic efficacy of two LAB strains is highly dependent on experimental obesogenic diets. These findings highlight the need to carefully consider the confounding impact of diet in order to improve both the reproducibility of preclinical probiotic studies and their clinical research translatability.

**Graphical abstract:** 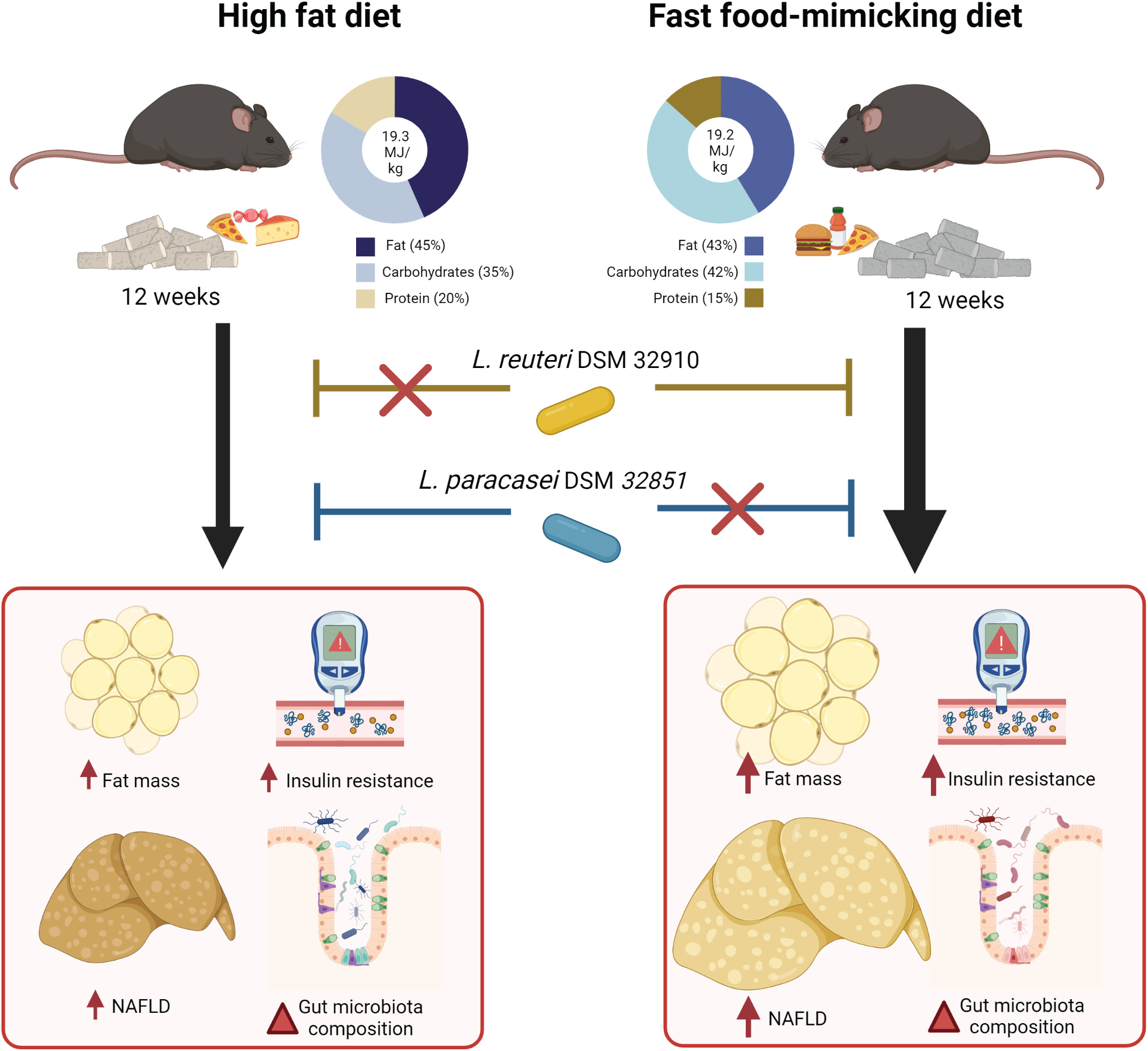

## Introduction

Since 1975 the global prevalence of overweight and obesity has nearly tripled to ~40% of the adult population ^1^. Obesity is a leading risk factor of cardiometabolic multimorbidity ^2^ including type 2 diabetes ^3,4^ and non-alcoholic fatty liver disease (NAFLD) ^5^, also known as metabolic associated fatty liver disease (MAFLD) ^6^, which affects ~25% of the population worldwide ^7^. The etiology of obesity is highly complex, involving positive energy balance resulting from poor dietary habits and sedentarity ^8^ but also features hereditary and socioeconomic traits ^9,10^. Although lifestyle interventions such as dietary restrictions may facilitate acute weight loss, this tends to decrease over time resulting in recurrent weight gain ^11–14^. Pharmacological therapies against obesity with proven clinical efficacy are emerging, however adverse effects such as nausea are frequently reported among users ^15^. Surgical strategies, most notably gastric bypass, are more successful yet highly invasive and not spared from potential perioperative complications ^16^.

Probiotics - defined as “live microorganisms which when administered in adequate amounts confer a health benefit on the host” ^17^ - exhibit promising preclinical potential to reduce obesity, NAFLD, and insulin resistance. Probiotics are therefore considered a non-invasive yet potentially highly effective therapeutic approach against obesity and associated metabolic diseases ^18,19^. Still, evidence from clinical trials is inconsistent and often of low quality, as the probiotic field is loosely regulated and evidence required for approval by authorities highly dependent on geographical region ^20,21^. This emphasizes a critical need for an enhanced molecular understanding of strain-specific capabilities ^22^ including potential adverse effects, as reported recently for probiotic use in cancer immunotherapy ^23^. Human studies with probiotic supplementation further reveal responders and non-responders as a common trait, both in terms of colonization and general host effects ^24,25^. Whether the lack of consistent probiotic efficacy in previous studies relates to methodological, genetic and/or environmental factors, such as dietary patterns, remains largely unknown ^24,25^.

A limited number of studies have reported efficacy discrepancies of both lactic acid bacteria (LAB) strains and other taxa in mice depending on the diet they consumed. Indeed, *L. helveticus* R0052 induced different effects on anxiety-like behavior depending on whether the mice consumed a fiber-rich chow or high-fat Western diet ^26^, which was also the case for *L. plantarum* WCFS1’s effects on bacterial gene expression and host intestinal inflammation ^27^. On a similar note, *Prevotella copri* appears to exhibit opposing effects on glucose metabolism in both humans and mice depending on dietary challenge, being either a fiber-rich low fat diet ^28^ or a low fiber, high fat diet ^29^. Importantly, these studies were confounded by dietary differences in energy and fiber content. To our knowledge, no studies have yet examined probiotic efficacy in different but energy-matched obesogenic diets resulting in distinct metabolic phenotypes and disease severity in preclinical murine models. This is critically important to optimize the clinical translatability of preclinical research in the probiotic field.

In this study, we therefore evaluated the efficacy of two LAB strains in mice fed either a standard HFD commonly used in the field or a customized diet designed to exacerbate NAFLD and nonalcoholic steatohepatitis (NASH). We demonstrate that the efficacy of the tested bacterial strains on preventing obesity, insulin resistance, and NAFLD development is entirely dependent on the obesogenic diet used, underlining the critical importance of considering the composition of experimental diets in preclinical studies for evaluating the effects of probiotics. Our work further highlights the need to carefully assess dietary patterns and nutritional factors as potential confounders of probiotic efficacy in clinical studies.

## Materials & methods

### Animals study design and diets

Three independent studies with identical experimental outlines were carried out using male C57BL6/J mice (Jackson Lab) co-housed three mice per cage. Twelve days prior to the start of the study period the mice were changed from a chow diet to a compositionally defined low fat diet (LFD) during acclimatization. Experimental groups were stratified based on similar average and variance of baseline body weight, fat mass, and additionally for Study 1, 5h fasting blood was sampled for determinations of plasma glucose and insulin levels before starting the experimental diets (described in a later section) without disturbing the social hierarchy within cages (**Figure S1A-E**). Study period was set to 12 weeks, allowing for significant weight gain, glucoregulatory disturbances and NAFLD onset ^30^. During the 12-week study period mice were fed experimental compositional defined diets (CDD) from eight weeks of age receiving either a commonly used high fat diet (HFD, D12451, provided by Ssniff Spezialdiäten (Study 1) or Research diets (Study 2), a customized fast food-mimicking diet (FFMD, D12079 mod. provided by Ssniff Spezialdiäten (Study 1 and 3) or continued on the LFD by Ssniff Spezialdiäten consisting of dietary sources matched to the FFMD (**Fig 1A, Table 1**). Feed intake was monitored and exchanged three times a week and body weight was measured weekly. Body composition was assessed by magnetic resonance (MR) scan (Minispec LF290 NMR analyzer, Bruker) prior to, and then every four weeks during the study. Energy uptake was measured at week 11 of the study by monitoring the feed intake and fecal excretion during 24h in clean cages with bedding followed by energy excretion using a calorimeter. Due to co-caging energy in- and uptake was only possible to assess at cage-level and presented as average/mouse/cage with n-size equalling the number of cages per group. From the start of feeding experimental diets, mice were orally gavaged daily 4h into the light cycle with 100 μL of either vehicle (PBS, Gibco) or 10^8^ CFU *Limosilactobacillus reuteri*, formerly known as *Lactobacillus reuteri*, DSM 32910 (*L. reuteri*) or *Lacticaseibacillus paracasei* subsp. *paracasei*, formerly classified as *Lactobacillus paracasei* subsp. *tolerans*, DSM 32851 (*L. paracasei*) freshly thawed from glycerol stocks stored at −80°C and the live cells diluted in PBS (Gibco). After twelve weeks, the mice were anaesthetized with isoflourane (Fresenius Kabi) and euthanized by cervical dislocation in alternating order after 5h of fasting from 2h into the light cycle. Cardiac puncture was performed using a 25G needle and 1 mL syringe coated with EDTA (Sigma-Aldrich). Blood was transferred to Eppendorf tubes containing 1 μL of DPP IV inhibitor (Millipore) and 1 μL of a protease inhibitor cocktail (Sigma) and kept on ice. The samples were centrifuged at 1000 rcf at 4 °C for 10 min and the plasma was placed on dry ice and transferred to −80 °C storage until further processing.

**Figure 1:**
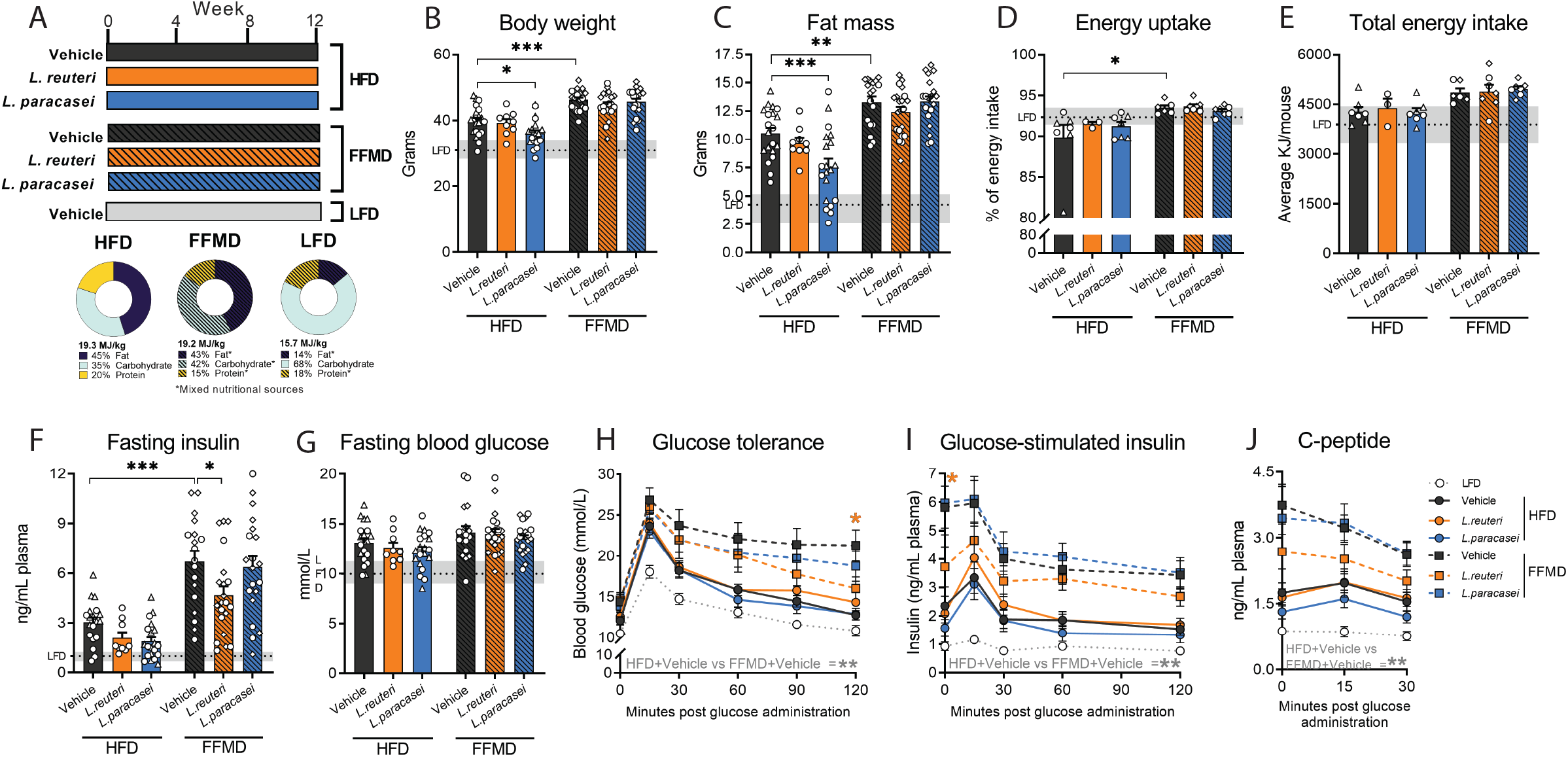
Body composition and insulin sensitivity are diet-dependently affected by probiotic strains. **A)** Experimental outline for all three studies with 12 weeks of feeding either High Fat Diet (HFD), Fast Food-mimicking Diet (FFMD) or Low Fat reference Diet (LFD) (matched to the FFMD in nutritional sources) with indicated total energy content in MJ/kg and energy distribution of macronutrients in percentage. Shades indicate mixed nutritional sources of the macronutrients. During the study, the mice were orally gavaged daily with either 10^8^ CFU of either *Limosilactobacillus reuteri* (*L. reuteri*), *Lacticaseibacillus paracasei* subsp. *paracasei* (*L. paracasei*), or an equal volume of vehicle (PBS). **B)** Body weight in grams at the end of the study period. **C)** Fat mass in grams assessed by MR scan at the end of the study period. **D)** Energy uptake per cage monitored over 24h as the inverse percentage of energy excretion compared to energy intake assessed in week 11 of the study. **E)** Accumulated energy intake per cage during the study period as average intake per mouse. **F)** Plasma 5h fasting insulin assessed in week 10 of the study. **G)** Blood glucose levels of 5h fasted mice in week 10 of the study. **H)** Blood glucose values from Study 1 during oral glucose tolerance test (oGTT) with 1.5 μg/g lean mass dextrose in week 10. **I)** Corresponding plasma insulin levels during oGTT. **J)** Corresponding plasma C-peptide levels during oGTT. **B-G)** Bars indicate group mean ± SEM. Points represent individual mice; individual studies are identified by point shapes where circles indicate Study 1, triangles Study 2, and squares Study 3. Dashed line indicates LFD group mean with interquartile range shown in grey. Asterisks indicate fdr-corrected q-values <0.05 using linear mixed effects models comparing indicated groups. **H-J)** Lines indicate group mean ± SEM. Asterisks indicate p-values <0.05 using two-way ANOVA with multiple comparisons between vehicle-treated groups or *Lactobacillus* treated group to vehicle-treated group fed the same diet with Bonferroni post-hoc test. **B, C, F, G)** LFD+Vehicle n=21, HFD+Vehicle n=20, HFD+L.reuteri n=9, HFD+L.paracasei n=20, FFMD+Vehicle n=18, FFMD+L.reuteri n=21, FFMD+L. paracasei n=20. **D-E)** LFD+Vehicle n=7, HFD+Vehicle n=7, HFD+L.reuteri n=9, HFD+L.paracasei n=7, FFMD+Vehicle n=6, FFMD+L.reuteri n=7, FFMD+L. paracasei n=7. **H-J)** LFD+Vehicle n=8, HFD+Vehicle n=9, HFD+L.reuteri n=9, HFD+L.paracasei n=9, FFMD+Vehicle n=9 (I-J n = 8 due to insufficient plasma volume from one mouse), FFMD+L.reuteri n=9, FFMD+L. paracasei n=8.

**Table 1:**
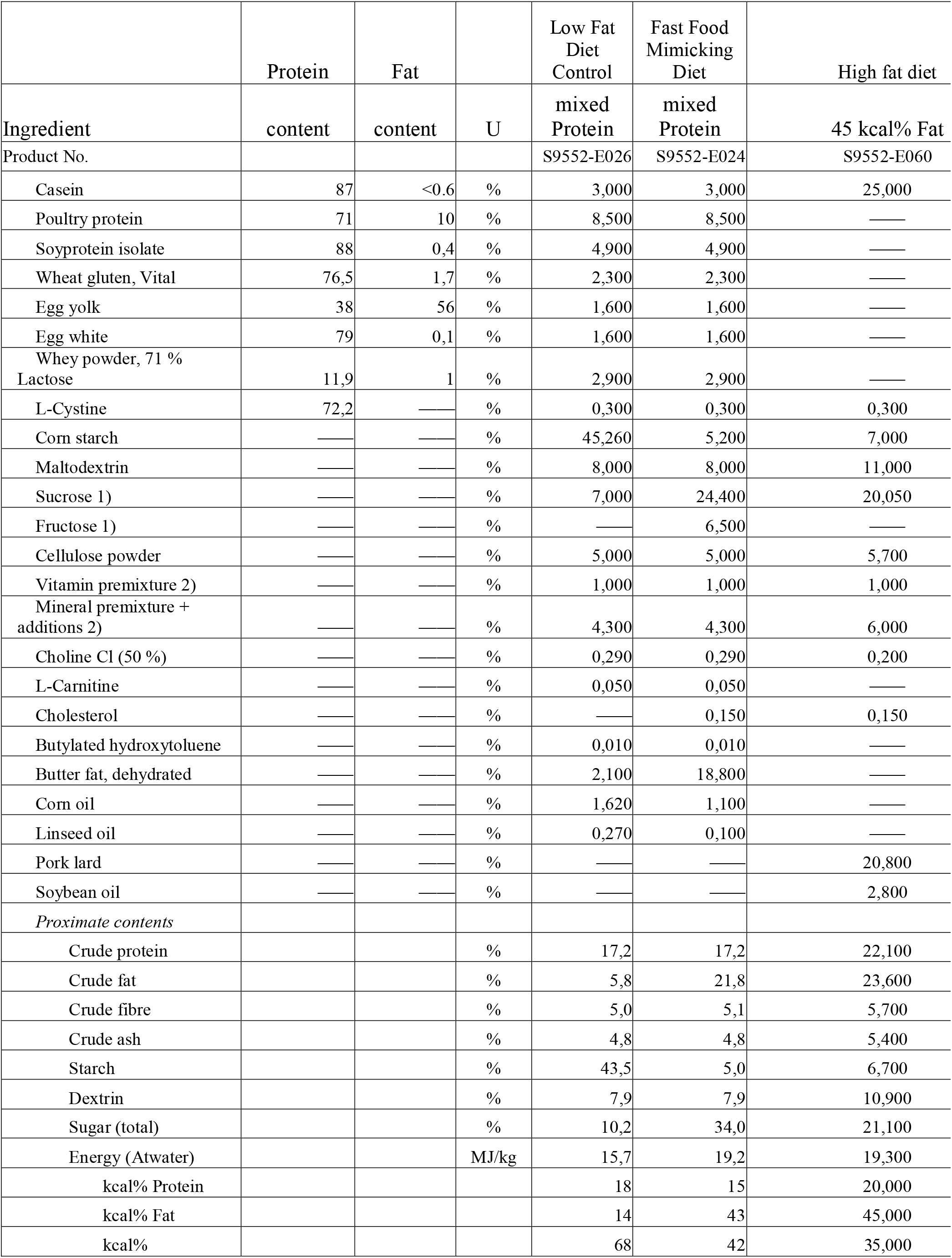
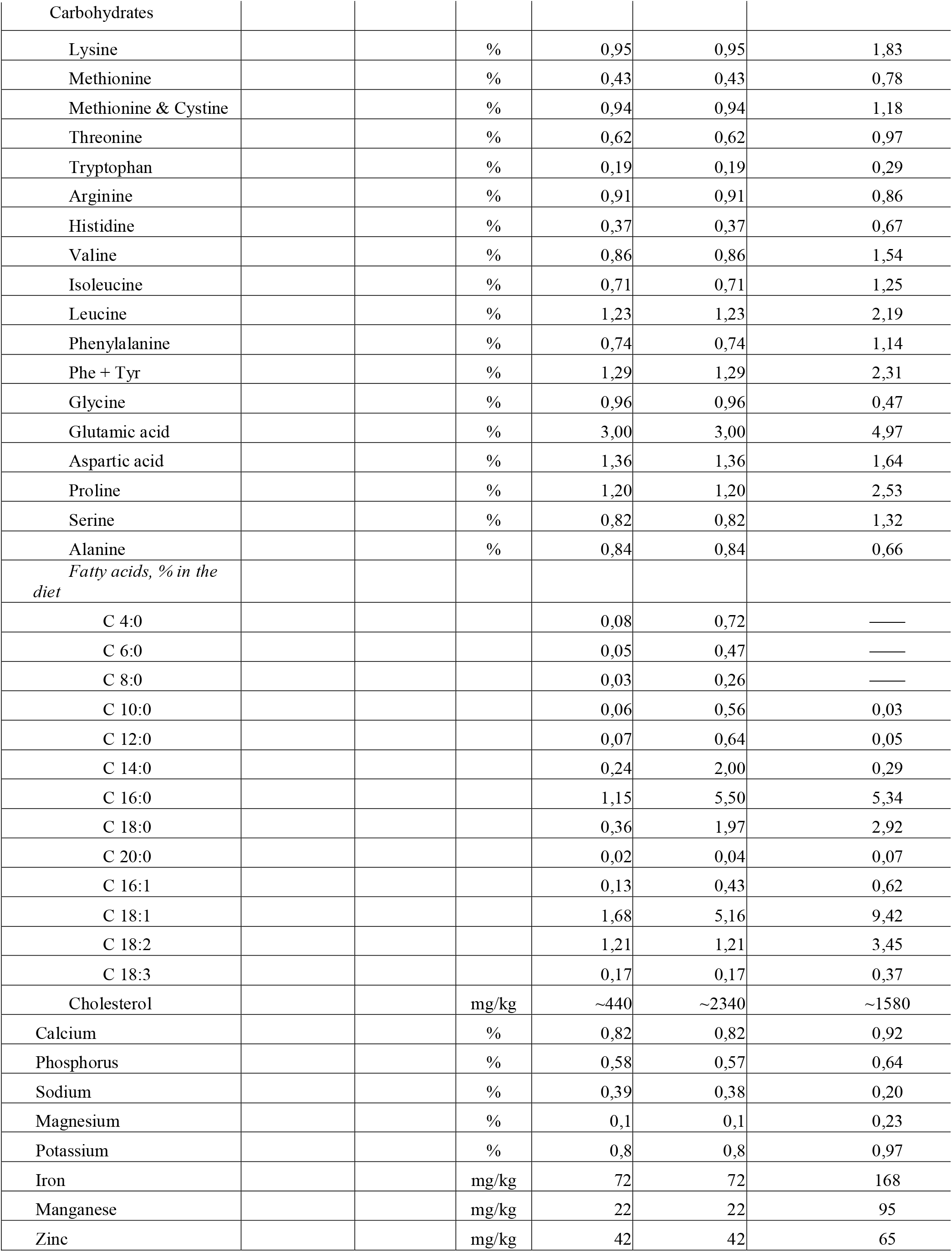
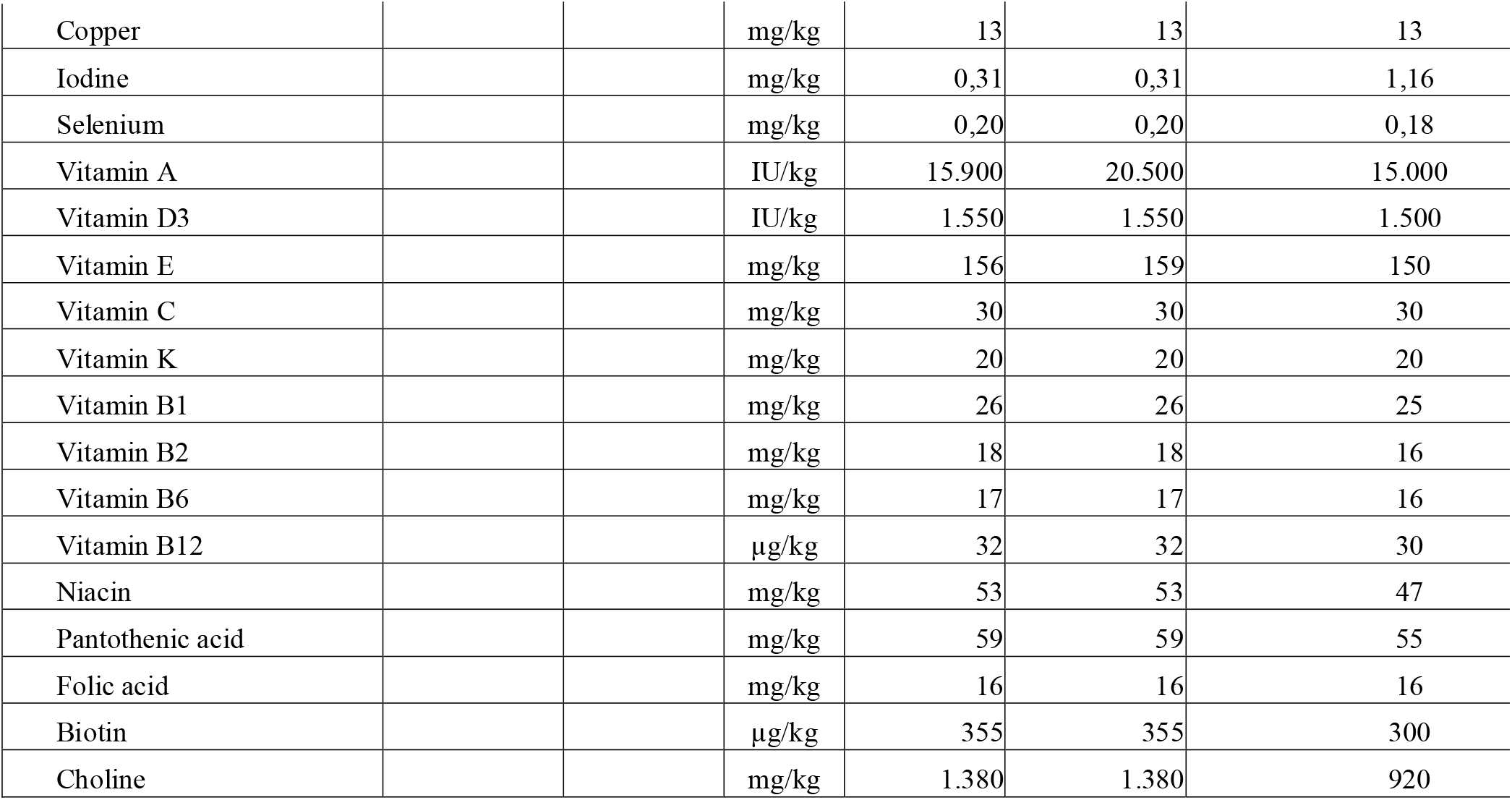
Composition of diets.

The weight of the liver, white adipose tissues (WATs), and cecum were measured, and the tissues immediately frozen in liquid nitrogen and stored at −80 °C until further processing. Tissues were dissected by the same operator and taken in the same order for all mice in each study. The small intestine (from stomach to cecum) and colon (from cecum to rectum) were kept on a Plexiglas plate cooled by underlying wet ice throughout the handling time. Duodenum was considered the first 5 cm of the small intestine and the remaining small intestine tissue was divided into 3 parts of equal length. The proximal 3 cm of the distal 1/3 of the small intestine was discarded and the remaining tissue categorized as ileum. Content of the small intestine, cecum, and colon were isolated by mechanical pressure, frozen on dry ice, and subsequently stored at −80 °C. Tissues from the ileum and colon were snap-frozen in liquid nitrogen and stored at −80 °C. The animal studies were conducted in accordance with Canadian Council on Animal Care under the Laval University license 2017-086.

### Assessment of glucose homeostasis, oral glucose tolerance, glucose-stimulated insulin, and C-peptide levels

Study 1 baseline glucose homeostasis was assessed in 5h fasted mice the day of study initiation. Blood glucose was measured by sampling blood from the tail vein using a glucometer (OneTouch Vario Flex, LifeScan) and plasma sampled using EDTA prepared capillary tubes (Sarstedt) kept on ice until blood samples were centrifuged for 10 min at 1000 rcf at 4 °C. Plasma insulin was quantified by Mouse Ultrasensitive Insulin ELISA (Alpco) using the manufacturer’s protocol.

An oral glucose tolerance test was carried out in week 10 of the experimental protocol. Mice were fasted from 2h into the light cycle for 5h still receiving their usual oral daily gavage. Fasting blood glucose and plasma insulin were measured as described for baseline glucose homeostasis prior to oral gavage with 1.5 μg/g (Study 1 and 3) or 2.0 μg/g (Study 2) lean mass of dextrose. Blood glucose was measured from tail vein puncture at time points 15, 30, 60, 90, and 120 min after dextrose challenge. Blood samples for quantification of plasma insulin levels were sampled in EDTA prepared capillary tubes (Sarstedt) at time points 0, 15, 30, 60, and 120 min and for C-peptide at time points 0, 15, and 30 min post oral glucose challenge. Mice were subcutaneously injected with 0.5 mL saline (Hospira) after the procedure for rehydration. Blood samples were centrifuged for 10 min at 1000 rcf at 4 °C. Plasma insulin was quantified by Mouse ultrasensitive Insulin ELISA (Alpco) and C-peptide by Mouse C-peptide ELISA kit (Crystal Chem) following the manufacturers’ protocols.

### Liver triacylglycerol (TG) quantification

TG was quantified in snap-frozen liver tissue stored at −80 □C until cryo-grinding in liquid nitrogen and 50 ±5 mg tissue added 0.9 mL of a 2:1 chloroform:methanol solution and homogenized 1 min at 50 os/sec using a TissueLyser LT (Qiagen) with beads. The content was transferred to new tubes and added 0.3 mL methanol and incubated for 2h with rotation. The tubes were centrifuged for 15 min at 4000 rcf at 10°C followed by transfer of 0.825 mL supernatant to new tubes and 0.4 mL chloroform was added. The samples were vortexed, 0.275 mL of 0.73% NaCl was added and samples were vortexed again for 30 sec followed by centrifugation for 3 min at 4000 rcf at 10°C. The upper phase was aspirated and the tubes rinsed with 0.8 mL Folch solution (Chloroform:Methanol:NaCl 0.58% (3:48:47)) repeatedly for three washes followed by addition of 2-5 drops of methanol until the solution appeared clear after vortexing. The liquid was evaporated using liquid nitrogen and the extracted lipids were resuspended in a solution of isopropanol with 10% TritonX-100 and TG levels quantified by Infinity Triglycerides Reagent (ThermoFisher) and calculated from serial diluted standard curves.

### Lipidomics of liver tissue and plasma

Liver and plasma lipids were extracted in a solvent consisting 89.9:10:0.1 of Heptan/IPA/HCOOH and analyzed by liquid chromatography-mass spectrometry (LC-MS) (Thermo Scientific). The MS was operated in negative and positive ion mode electrospray ionization mode. Data was collected from 80-1500 m/z. Lipids were normalized to sample median by log transformation and Pareto scaling using MetaboAnalyst ^31^, which was additionally used to generate heatmaps of the top 40 or 100 features of each measure as indicated in figure legends.

### Histopathological scoring of liver tissue

Liver tissue (lobus dexter medialis hepatis) was collected and fixed in 4% paraformaldehyde for three days. After paraffin embedding 3 μm sections were prepared and stained with hematoxylin and eosin (H&E) or Sirius Red. NAFLD activity was assessed blindly in H&E-stained cross-sections using an established scoring system for murine NAFLD ^32^. In short, the levels of hepatocellular hypertrophy, macrovesicular and microvesicular steatosis were determined at 40× to 100× magnification relative to the total liver area and scored as described in **Supplementary Table 1**. Hepatic inflammation was scored by counting the number of inflammatory foci per 1 mm^2^ using a 100× magnification. Inflammatory foci were defined as clusters (not rows) of at least 5 inflammatory cells and the mean of five random fields was used for scoring, as previously described ^32^. Histopathological categories were based on previously described methodology ^32^ where liver slides are categorized with NAFLD if the steatosis score (based on the sum of micro-, macrovesicular steatosis, and hepatocellular hypertrophy scores) ≥ 1 with an inflammation score of 0. Mice are categorized with NASH if steatosis score ≥ 1 and an inflammation score ≥ 1. Livers are considered healthy if the steatosis score = 0 regardless of the inflammation score. Fibrosis was blindly assessed by the Ishak fibrosis stage using Sirius Red-stained cross-sections following previously published methods ^33^. The scoring system introduced by Ishak *et al*. allows for a sensitive scoring of fibrosis that takes quantity and location of fibrosis into account ^34^ and is described further in **Supplementary Table 2**.

### Cytokine quantification in liver and adipose tissue

Cryo-grinded liver tissues were washed by adding 200 μL ice-cold PBS and protease inhibitors (P8340 Sigma-Aldrich), centrifugation at 16,000 x g for 20 sec at 4 °C and aspiration of the liquidphase. This wash step was repeated a total of 10 times. Afterwards 400 μL T-PER Tissue Protein Extraction Buffer (ThermoFisher) and beads were added and vortexed for 2 x 1 min at 50 os/sec. Samples were centrifuged at 4,000 rcf for 1 min at 4 °C and the supernatants transferred to new tubes that were centrifuged again at 13,000 rcf for 10 min at 4 °C. Snap-frozen mWAT was broken into pieces of 100-200 mg tissue then added acid washed zirconium beads (OPS Diagnostics) while kept frozen. mWAT samples were homogenized in ice-cold lysis buffer (PBS + 1% IGEPAL CA-630 (Sigma) plus protease inhibitors (P8340 Sigma-Alrich)) added 500 μL/100 mg tissue using a Bead Mill homogenizer (VWR) and continuously rotated at 4 °C for 30 min. Samples were centrifuged at 12,000 rcf for 15 min at 4 °C, supernatants transferred to clean tubes to repeat the centrifugation for 10 min. Protein concentration from liver and mWAT extracts were measured in triplicates using Pierce BCA Protein Assay kit (ThermoFisher) following the manufacturer’s instructions. Cytokine levels in samples normalized to the protein concentration were measured using ProcartaPlex Immunoassay (ThermoFisher) for liver samples or Bio-Plex Pro Mouse Th17 Cytokine Assay (Bio-Rad) for mWAT samples following the manufacturers’ instructions on the Bio-Plex 200 system (Bio-Rad).

### Bile acid (BA) characterization in liver tissue and cecum content

BAs in liver tissue and cecum content were extracted with methanol and 5 μL was injected onto a reverse-phased chromatography (1290 Infinity II, Agilent) and mass spectrometer (6546 Q-TOF/MS, Agilent). The MS was operated in negative ion mode electrospray ionization. Data were collected from 80-1600 m/z. The chromatographic part of the LC-MS-system was set up with a CSH-peptide C18 column (Waters) using a gradient system with Eluent A: water with 0.1% formic acid and Eluent B: Acetonitrile with 0.1% formic acid. A flow of 0.250 mL/min was used, starting at 99.5% Eluent A lowering to 1% within 19 min followed by a new gradient endpoint 99.5% Eluent A at 25.5 min. The gradient was kept isocratic until 30 min and the eluent returned to initial conditions at 26 min. All gradients were linear. Results were related to serial diluted standard curves.

### Tissue gene expression by quantitative reverse transcription PCR (RT-qPCR)

RNA from snap-frozen ileum and liver tissues were extracted using the Quick-RNA Miniprep Plus kit (Zymo Research). cDNA was synthesized from 1.5 μg ileum or 2 μg liver RNA using High-Capacity cDNA Reverse Transcriptase kit (Applied Bioscience) following the manufacturer’s protocol. qPCR was carried out using 4μl of cDNA, 5μl of Advanced qPCR MasterMix (Wisent Bioproducts) and 0.5μl of each primer (diluted at a concentration of 10 μM) in a total reaction volume of 10 μL with the following cycle setting: 95 °C for 2 minutes, (95 °C for 20 sec, 61.5 °C for 20 sec, 72 °C for 20 sec) x 40 ending with a melting curve: 65 °C to 95 °C. The targets were evaluated in each tissue and accepted in case of a unified peak from the melting curve and amplification efficiency of 100%±11 and R^2^ > 0.99 from a standard curve. Relative expression was calculated by 2^ΔCq^ of target Cq to 18S Cq of the sample accepting replicates with coefficient of variation < 0.02. Target primer sequences are reported in **Supplementary Table 3**.

### Microbiota profiling by 16S rRNA gene amplicon sequencing

Fecal samples were collected prior to the start of the studies and every 4 weeks throughout the study period at the same time as body composition assessment 22-24h after the latest oral gavage with probiotic strains or vehicle. During necropsy 2-5h after the latest oral gavage contents of small intestine and colon were collected. All samples were immediately placed on dry ice and stored at −80 °C until DNA extraction using the NucleoSpin 96 soil kit (Macherey-Nagel) following the manufacturer’s instructions. Stocks of the administered LAB strains were processed similarly to the remaining samples. The 16S rRNA gene was amplified over 25 cycles using primers for the V3-V4 region with Illumina adaptors (S-D-Bact-0341-b-S-17: *5’-TCGTCGGCAGCGTCAGATGTGTATAAGAGACAGCCTACGGGNGGCWGCAG-3’and* S-D-Bact-0785-a-A-21: *5’-GTCTCGTGGGCTCGGAGATGTGTATAAGAGACAGGACTACHVGGGTATCTAATCC-3’*)^35^. Indices were added in a second PCR over 8 cycles with unique primer combinations using the Nextera XT Index Kit V2 (Illumina). The samples were pooled and cleaned using AMPure XP beads (Beckman Coulter) and the library was sequenced on an Illumina MiSeq desktop sequencer using the MiSeq Reagent Kit V3 (Illumina) for 2x 300 bp paired-end sequencing. The generation of an amplicon sequence variant (ASV) table was carried out using usearch version 10.0.240. Primer binding regions were removed with fastx_truncate and reads were filtered to contain less than one error per read. The quality filtered reads were denoised with unoise3. ASV abundance was calculated by mapping with usearch global using a 97% identity threshold. The phylogenetic tree was made by aligning the 16S sequences with mafft, and the tree was inferred by FastTree. Taxonomical classification was done with the qiime classifier (qiime2-2019.4) trained on the Silva database (Silva_132) as previously described ^36^. The dataset had a median of 45,468, a mean of 46,400 reads per sample with a standard deviation of 19,177 and including samples with a minimum of 10,000 reads. Alpha diversity measures and analysis of differential abundances were calculated based on rarefied data. Principal coordinate analysis (PCoA) of microbiota composition was carried out using weighted UniFrac distances. Functions of the microbiota were predicted using the Kyoto Encyclopedia of Genes and Genomes (KEGG) orthology generated by Picrust2 and illustrated with PCoAs using Bray-Curtis distances. Specific changes in regulated predicted pathways were tested using DeSeq2 between groups of interest reporting p-value and false discovery (fdr)-corrected q-values.

### Correcting for multiple hypothesis testing

For all analyses described below and throughout the manuscript, nominal p-values were corrected for multiple hypothesis testing to fdr (q-values) using the Benjamini-Hochberg method ^37^.

### Software and Statistical analysis

The statistical analysis is specified in the figure legend of each figure. Statistical analysis by linear mixed-effects models was carried out in R using the lmerTest package ^38^ including study, batch, and cage as nested random effects. Statistical significance was illustrated by asterisks based on fdr-corrected q-values. Two-way analysis of variance (ANOVA) were conducted to analyze the effects of treatments and diet in addition to Bonferroni multiple comparison test to respective vehicle-treated control on the same diet. Categorical data was tested for statistical differences by Kruskal-Wallis and Mann-Whitney tests with Dunn’s post hoc for multiple comparisons testing. Statistical analysis of lipidomics data was carried out by one-way ANOVA with Tukey’s post-hoc analysis in the MetaboAnalyst software. The graphical abstract was created with BioRender.com. Bar charts were made using GraphPad Prism 9. Analysis of community-based statistical differences on gut microbiota composition and its predicted functions were carried out by permutational multivariate analysis of variance (PERMANOVA) by the R package vegan ^39^ using weighted UniFrac or Bray-Curtis distances, respectively, reporting an average p-value of 100 permutations. The glmer.nb function from the lme4 package ^40^ fitting a generalized linear mixed-effects model (GLMM) for the negative binomial family building on glmer was used for the analysis of differential abundance of bacterial phylae on rarefied data. The top 100 most abundant taxa on each taxonomy rank were used in the analysis. Cage was used as random effect in the model. Multiple testing correction was done with the fdr method. To provide full transparency of reported findings, parameters assessed in independent animal studies were shown in the same graph with experiment-dependent point shapes, as indicated in figure legends when appropriate, allowing the reader to visually assess experiment-to-experiment variation. To avoid unwarranted power inflations, we used linear mixed-effects model as described above, including study, batch, and cages as nested random effects and only reports fdr-corrected q-values. Collectively, these precautions and strict statistical analysis enhance experimental reproducibility, and thus study validity. Study 1 sample size (n) = 9 mice per group except for FFMD + *L. paracasei* group with n = 8. Study 2 n = 11 mice per group. Study 3 n = 12 mice per group except for FFMD + Vehicle group with n = 9. Asterisks identify statistically significant fdr-corrected q-values where * = < 0.05, ** = < 0.01 and *** = < 0.001.

## Results

### Experimental diet determines probiotic efficacy on body composition and insulin sensitivity

We ensured equal starting points between the experimental groups and fed the mice one of two energy-matched high fat, high sucrose diets or a low fat reference diet (LFD) for 12 weeks with daily oral gavage of either *L. reuteri, L. paracasei*, or vehicle (**Fig 1A**, **S1A-E, Table 1**). Despite similar feed energy content, vehicle-treated FFMD-fed mice exhibited enhanced body weight-, fat- and lean mass gain compared to their classical HFD-fed counterparts (**Fig 1B-C, Fig S1F**). This trait was closely associated with an increase in energy uptake (**Fig 1D**) and a similar trend for energy intake (p = 0.02, q = 0.11, **Fig 1E**). Of note, fat percentage, subcutaneous, epididymal and retroperitoneal white adipose tissues (WATs) were equally increased by FFMD and HFD, whereas FFMD augmented visceral mesenteric WAT deposition to a higher degree than HFD resulting in higher total mWAT inflammation as revealed by the increased levels of several pro-inflammatory cytokines in the tissue, which were otherwise not significantly influenced by probiotic treatments (**Fig S1G-N)**. *L. paracasei* reduced body weight and fat mass gain in HFD-, but not in FFMD-fed mice (**Fig 1B-C, S1G**). This effect of *L. paracasei* on body composition was not explained by a reduction of energy intake or uptake (**Fig 1D-E**).

We next investigated the effect of the the probiotic strains on glucose homeostasis after 10 weeks of feeding the obesogenic diets. Importantly, while the classical HFD impaired glucose homeostasis and increased both blood glucose and glucose-stimulated insulin secretion (GSIS) during glucose challenge compared to LFD-fed mice, FFMD-feeding significantly aggravated all of the above (**Fig 1F-J**). Strikingly, FFMD-fed mice more than doubled their fasting insulin levels compared to HFD-fed counterparts (**Fig 1F**). Moreover, although fasting blood glucose was unaffected by the type of obesogenic diet, FFMD-fed mice failed to return to baseline blood glucose levels two hours post challenge accompanied by a 2-3 fold increase in GSIS, suggesting an early onset of frank diabetes in the latter dietary model (**Fig 1G-J**). Notably, *L. reuteri* lowered fasting insulin levels and improved glucose tolerance, with a concomitant trend towards diminished GSIS based on both insulin and c-peptide excursions exclusively in FFMD-fed mice (**Fig 1F-J**, **S1M-N**). In contrast, *L. paracasei* improved glucose tolerance in HFD-fed mice after high dose (**Fig S1O-P**) but not low dose (**Fig 1H-J**) glucose challenge. Collectively, these data demonstrates that the metabolic benefits of two *Lactobacilli* strains are highly dependent on the energy-dense obesogenic diets used.

### NAFLD development was diet-dependently mitigated by probiotic strains

As a major glucoregulatory organ, we next assessed hepatic lipid accumulation. Both liver weight *per se* and TG concentration were >3-fold higher in FFMD-fed mice compared to their HFD-fed counterparts (**Fig 2A-C**). In agreement with enhanced liver TGs, FFMD feeding altered the hepatic lipid profile (**Fig S2A**), and increased the clinical marker of liver damage, plasma ALT, associating with a borderline reduction in AST/ALT ratio (p = 0.02, q = 0.09, **Fig 2D-E, S2C**). FFMD aggravated both histological prevalence and severity of diet-induced NAFLD, with ~90 % of the animals in the FFMD + Vehicle group having histopathologically-validated NASH, contrasting with only ~45 % of NASH observed in HFD + Vehicle mice (**Fig 2F-G**). Accordingly, FFMD-fed mice developed significantly increased hepatocellular hypertrophy and microvesicular steatosis, as well as excessive hepatic inflammation and moderate fibrosis (**Fig 2H-N, S2D-E**) all pointing towards aggravated liver damage ^41^ as compared to their HFD-fed counterparts. Of note, and in line with previous publications demonstrating that reference CDD stimulate modest fat accretion ^42,43^, ~5 % of the LFD-fed mice exhibited mild NAFLD (**Fig 2G**) possibly linked to incorporation of sucrose in the LFD diet to match the nutrient composition of obesogenic diets.

**Figure 2:**
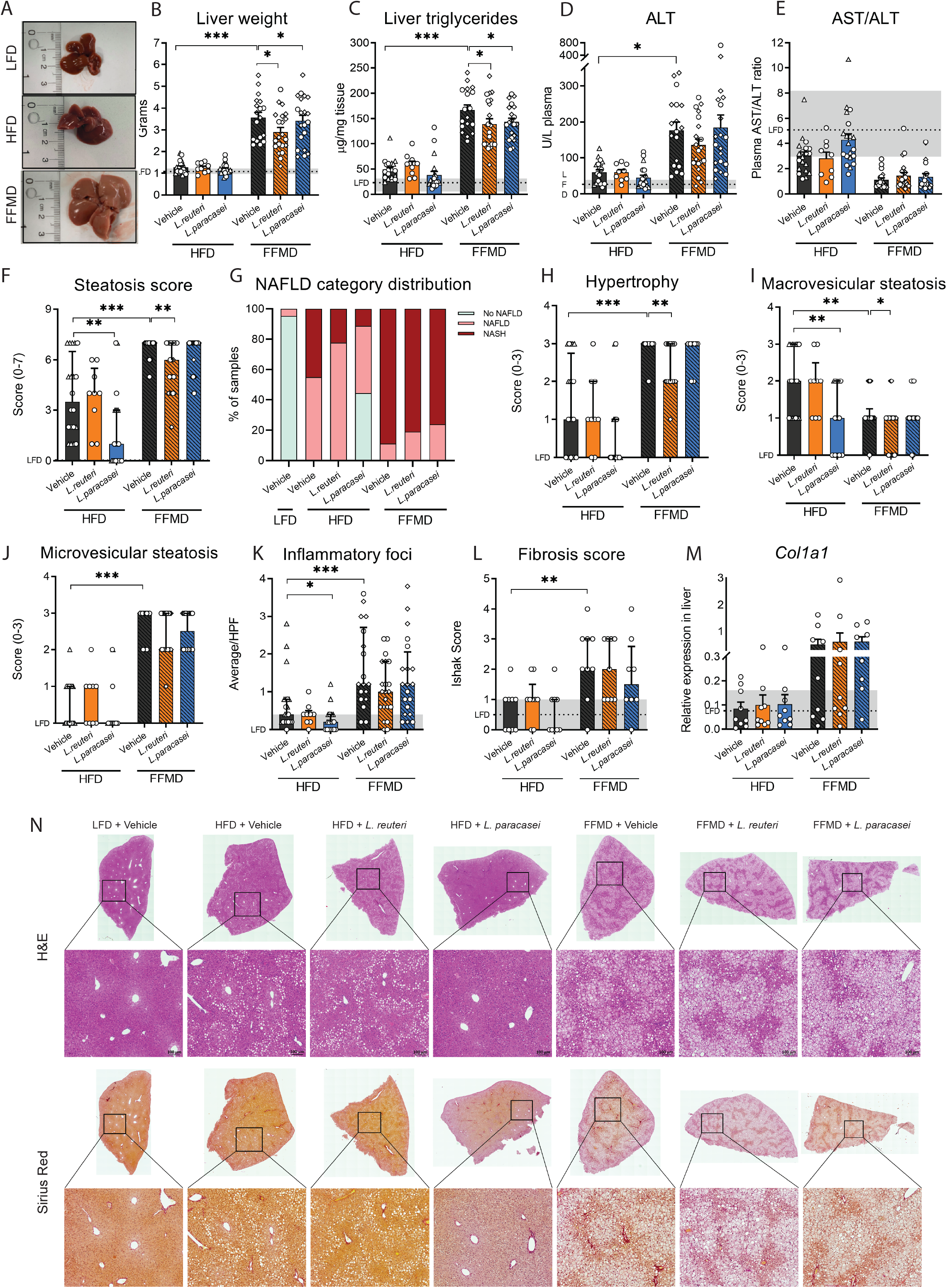
Non-alcoholic fatty liver disease (NAFLD) mitigation by probiotic strains depends on diet. **A)** Images of total livers from necropsy from representative mice in vehicle-gavaged dietary groups of Study 1 weighting respectively 1.01 g (LFD), 1.58 g (HFD) and 3.37 g (FFMD). **B)** Liver weight in grams at the end of the study. **C)** Liver triacylglyceride concentration as μg/mg tissue. **D)** Alanine transaminase (ALT) levels in plasma at the end of the study. **E)** Aspartate transaminase (AST) to ALT ratio in plasma. **F)** Histological steatosis score from 0-7 blindly assessed in H&E stained liver slides. **G)** Histopathological NAFLD categories as relative distribution per group of: No NAFLD, NAFLD, and NASH based on steatosis and inflammation scores of H&E stained liver slides. **H)** Hepatocellular hypertrophy score from 0-3 blindly assessed in H&E-stained liver slides. **I)** Macrovesicular steatosis score from 0-3 blindly assessed in H&E-stained liver slides. **J)** Microvesicular steatosis score from 0-3 blindly assessed in H&E-stained liver slides. **K)** Average number of inflammatory foci in 5 high power fields (HPF). **L)** Histological fibrosis assessed by Ishak scoring of Sirius Red-stained liver slides from Study 1. **M)** Relative gene expression of collagen type I alpha I chain (*Col1A1*) in liver tissue by qPCR. **N)** H&E and Sirius Red stained liver slides from Study 1 selected as representative by having the median value of the group in the histological steatosis score. **B-F, H-K)** Point shape represent individual mice in three individual studies where circles indicate Study 1, triangles Study 2, squares Study 3. **B-F, K-M)** Dashed line indicates mean of LFD group with interquartile range shown in grey. **B-E, M)** Bars indicate group mean ± SEM. Asterisks indicate fdr-corrected q-values <0.05 using linear mixed effects models comparing indicated groups. **F, H-L)** Bars indicate group median with interquartile range. Asterisks indicate p-values <0.05 by Kruskal-Wallis test with multiple comparisons between vehicle-treated groups or LAB group to vehicle-treated group fed the same diet with Dunn’s post-hoc test. **B-E)** LFD+Vehicle n=21, HFD+Vehicle n=20, HFD+L.reuteri n=9, HFD+L.paracasei n=20, FFMD+Vehicle n=18, FFMD+L.reuteri n=21, FFMD+L. paracasei n=20. **F-K)** LFD+Vehicle n=21, HFD+Vehicle n=20, HFD+L.reuteri n=9, HFD+L. paracasei n=20, FFMD+Vehicle n=20, FFMD+L. reuteri n=21, FFMD+L. paracasei n=20. **L-M)** LFD+Vehicle n=8, HFD+Vehicle n=9, HFD+L.reuteri n=9, HFD+L.paracasei n=9, FFMD+Vehicle n=9, FFMD+L.reuteri n=9, FFMD+L. paracasei n=8.

*L. paracasei* prevented diet-induced hepatic lipid accumulation in 17 out of 21 HFD-fed mice, presenting TG levels similar to those of LFD-fed reference mice (**Fig 2C**). A similar trajectory was noted for the hepatic lipid profile, where a subset of the HFD + *L. paracasei* group clustered together with their LFD-fed counterparts (**Fig S2B**). Additionally, the plasma AST/ALT ratio approached a normalization in HFD-fed mice receiving *L. paracasei* compared to the vehicle control (**Fig 2E**, p = 0.04, q = 0.09). The hepatic lipid clustering of the HFD + *L. paracasei* group fully recapitulated the histological steatosis score and NAFLD distribution (**Fig 2F-G, S2B**). Thus, HFD + *L. paracasei* mice clustering together with LFD-fed reference mice had a blinded steatosis score of 0-1, whereas those clustering together with HFD + vehicle control mice obtained a steatosis score ≥4, hence corroborating that NAFLD severity and not dietary feeding dictated hepatic lipid profile (**Fig 2F-G, S2B**). Both *L. reuteri* and *L. paracasei*, on the other hand, partly prevented the massively increased liver tissue mass and TG concentration in FFMD-fed mice (**Fig 2B-C**). When assessing the histological steatosis score, we observed that *L. paracasei* reduced steatosis development exclusively in HFD-fed mice, while only *L. reuteri* was able to significantly reduce this phenotype in FFMD-fed mice (**Fig 2F**). The selective NAFLD-reducing effect of *L. paracasei* in HFD-fed animals was reflected by a strong trend of diminished hepatocellular hypertrophy (p = 0.06) and macrovesicular steatosis (**Fig 2H-I, N**) hence mirroring the observed hepatic TG concentrations (**Fig 2C**). The selective amelioration of NAFLD by *L. reuteri* in FFMD-fed mice was mainly associated with reduced hepatocellular hypertrophy and a partial improvement of microvesicular steatosis (**Fig 2H, J**). *L. paracasei* reduced inflammatory foci counts in HFD-fed mice supported by a similar tendency in hepatic cytokine levels (**Fig 2K, S2D-E,** PERMANOVA p = 0.089), pinpointing the beneficial NAFLD ameliorating capabilities of *L. paracasei* in HFD-, but not FFMD-fed mice.

### *L. reuteri* diet-dependently altered intestinal bile acid levels

HFD and FFMD feeding distinctively impacted the plasma lipid profile (**Fig 3A**), where a majority of the altered lipids were regulated similarly to that observed in the liver (**Fig S2A**). Circulating levels of total cholesterol (**Fig 3B**) and TG (**Fig S3A**, p = 0.01, q = 0.06) were increased by FFMD compared to HFD while total free fatty acids (FA) were not significantly altered by the background diet (**Fig S3B**). A subset of the HFD-fed mice receiving *L. paracasei* clustered with the LFD reference group (**Fig 3A**), as seen with hepatic lipids (**Fig S2B)**. *L. reuteri* significantly increased the relative abundance of FA 18:2 exclusively in FFMD-fed mice (**Fig 3C**).

**Figure 3:**
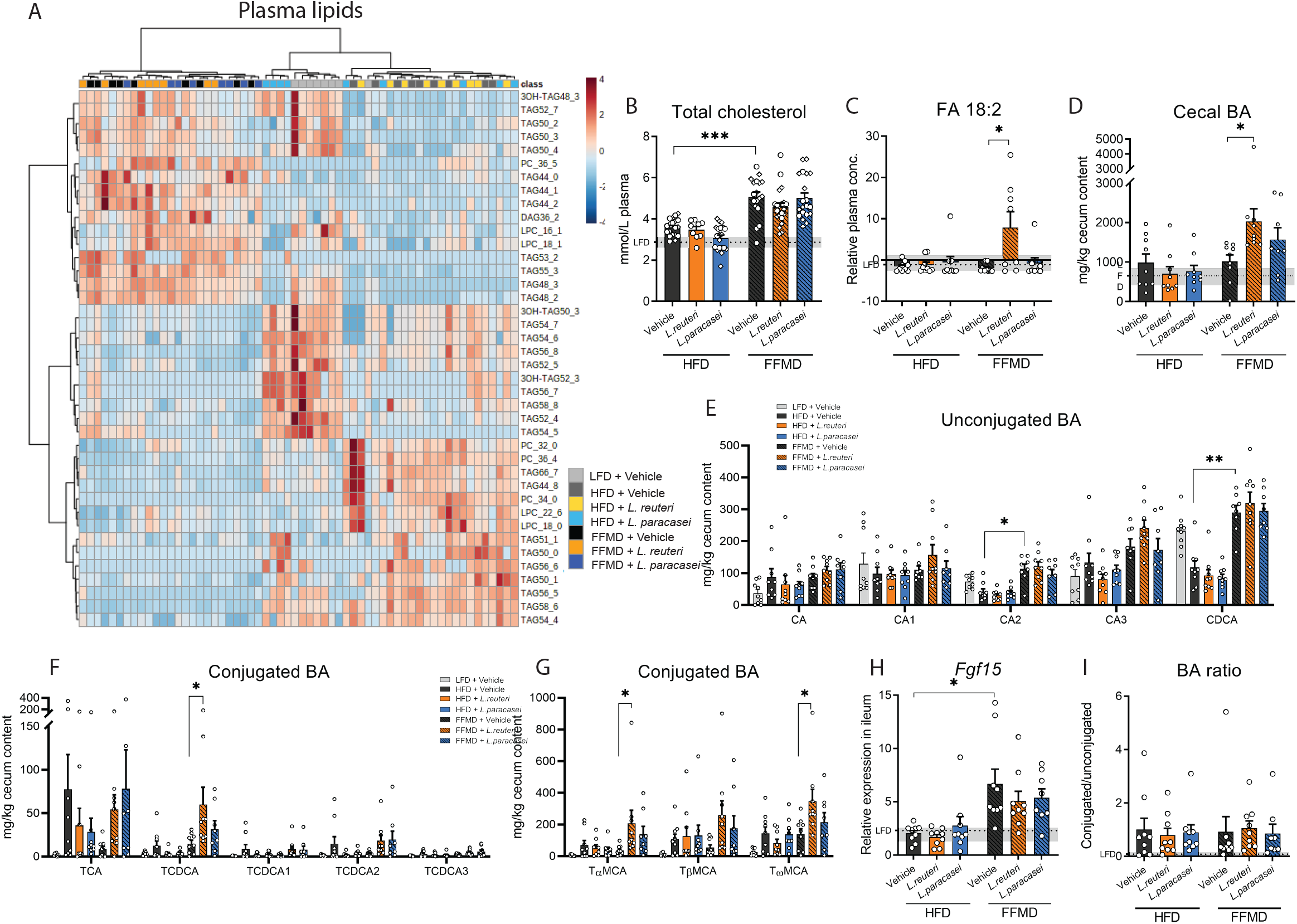
*L. paracasei* and *L.reuteri* diet-dependently affect liver lipids and cecum bile acids (BAs) **A)** Heatmap of the 40 most discriminating plasma lipids from end of study 1 assesed by lipidomics. Experimental groups are indicated by color and samples are distributed by hierical clustering. **B)** Total cholesterol levels in plasma. Point shape represent individual mice in three individual studies where circles indicate Study 1, triangles Study 2, squares Study 3. LFD+Vehicle n=21, HFD+Vehicle n=19, HFD+L.reuteri n=9, HFD+L.paracasei n=20, FFMD+Vehicle n=18, FFMD+L.reuteri n=21, FFMD+L. paracasei n=20. **C)** Relative plasma levels of free fatty acid (FA) 18:2 measured by lipidomics. **D)** Total measured BA concentration in cecum content. **E)** Concentration of unconjugated BA in cecum content as fold change to the LFD group. **F)** Concentration of conjugated cholic and chenodeoxycholic acids (CA and CDCA) in cecum content as fold change to the LFD group. **G)** Concentration of conjugated muricholic acids (MCA) in cecum content as fold change to the LFD group. **H)** Relative Fibroblast growth factor 15 (Fgf15) gene expression in ileum tissue by RT-qPCR. **I)** Ratio of measured conjugated to unconjugated BAs in cecum content. **B-I)** Bars indicate group mean ± SEM. Points represent individual data points. Asterisks indicate fdr-corrected q-values <0.05 using linear mixed effects models comparing indicated groups. **B-D, H-I)** Dashed line indicate mean of LFD group with interquartile range shown in grey. **C-I)** LFD+Vehicle n=9 (except C where n=8), HFD+Vehicle n=9, HFD+L.reuteri n=9, HFD+L.paracasei n=9, FFMD+Vehicle n=9 (except E where n=8), FFMD+L.reuteri n=9 (except C where n=8), FFMD+L. paracasei n=8.

Due to the known influence of bile acids (BAs) on the gut-liver axis by the enterohepatic pathway we quantified specific BAs in liver tissue. *L. paracasei* tended to increase total liver BA levels in HFD-fed mice (p = 0.02, q = 0.11) and the BA precursor THCA (**Fig S3C-D**) associated with borderline higher levels of an unconjugated and and taurine-conjugated BAs (**Fig S3E-F**, p < 0.05, q < 0.1). Over 90% of BAs excreted into the intestine are recirculated in the small intestine ^44^ prompting us to quantify the remaining BAs in the cecum, which did not differ in weight between the groups (**Fig S3G**). FFMD increased the levels of unconjugated BAs in this intestinal segment compared to HFD as well as secondary BAs without affecting the total cecal BA concentration (**Fig 3D-E, S3H**). These increases were associated with higher expression of fibroblast growth factor (*Fgf*)15 in ileum tissue (**Fig 3G**), which is upregulated by BAs through intestinal Fxr ^44^. The indication of enhanced activation of the Fxr-Fgf15 enterohepatic pathway did, however, not result in repression of *Cyp7a1* expression (**Fig S3I**), the rate-limiting enzyme of BA synthesis in the liver ^44^ thus suggesting a dysregulation of the pathway by FFMD feeding. The probiotics did not significantly alter *Fgf15* or host-defense protein *RegIIIγ* gene expression in ileum tissue on the respective diets (**Fig 3H, S3J**).

Despite the tendency of augmented hepatic BA levels in *L. paracasei* treated HFD-fed mice (**Fig S3C-F**), we did not observe changes in cecal BA levels in this group (**Fig 3E-G**). Conversely, *L. reuteri* increased cecal (**Fig 3D**) but not hepatic (**Fig S3C**) BA levels exclusively in FFMD-fed mice. Notably, only absolute abundance of conjugated BAs were affected by *L. reuteri* (**Fig 3E-G, I**), suggesting a reduced re-uptake of the conjugated BA as proposed for other *L. reuteri* strains ^45,46^ although this was not mirrored by altered gene expression of BA transporter *Asbt* in ileum tissue (**Fig S3K**). Investigation of the strains’ genomic makeup revealed that *L. reuteri*, but not *L. paracasei*, habored the bile salt hydrolase (*Bsh*) gene (PFAM PF02275), potentially driving the changes in intestinal BA content in a diet-dependent manner.

### Predicted functions of the gut microbiota correlated with disease severity, not macronutrient composition, and were modulated by probiotics

We collected fecal samples at weeks 0, 4, 8, and 12 to assess longitudinal changes in gut microbiota composition as a function of intervention and diet. At termination, we further collected the content of small intestine and colon. Microbiota composition was markedly affected by dietary macronutrient composition (FFMD and LFD clustering separately from HFD), in colonic and fecal samples, suggesting that dietary nutrients, such as proteins, carbohydrate or lipid sources, rather than lipid proportion or the occurrence of obesity determined gut microbiota composition (**Table 1, Fig 4A-C, S4A-C**). This was despite equal baseline microbiota composition and with a transient increase of fecal alpha diversity measures from FFMD and LFD (**Fig S4A, E-H, Table 2**).

**Figure 4:**
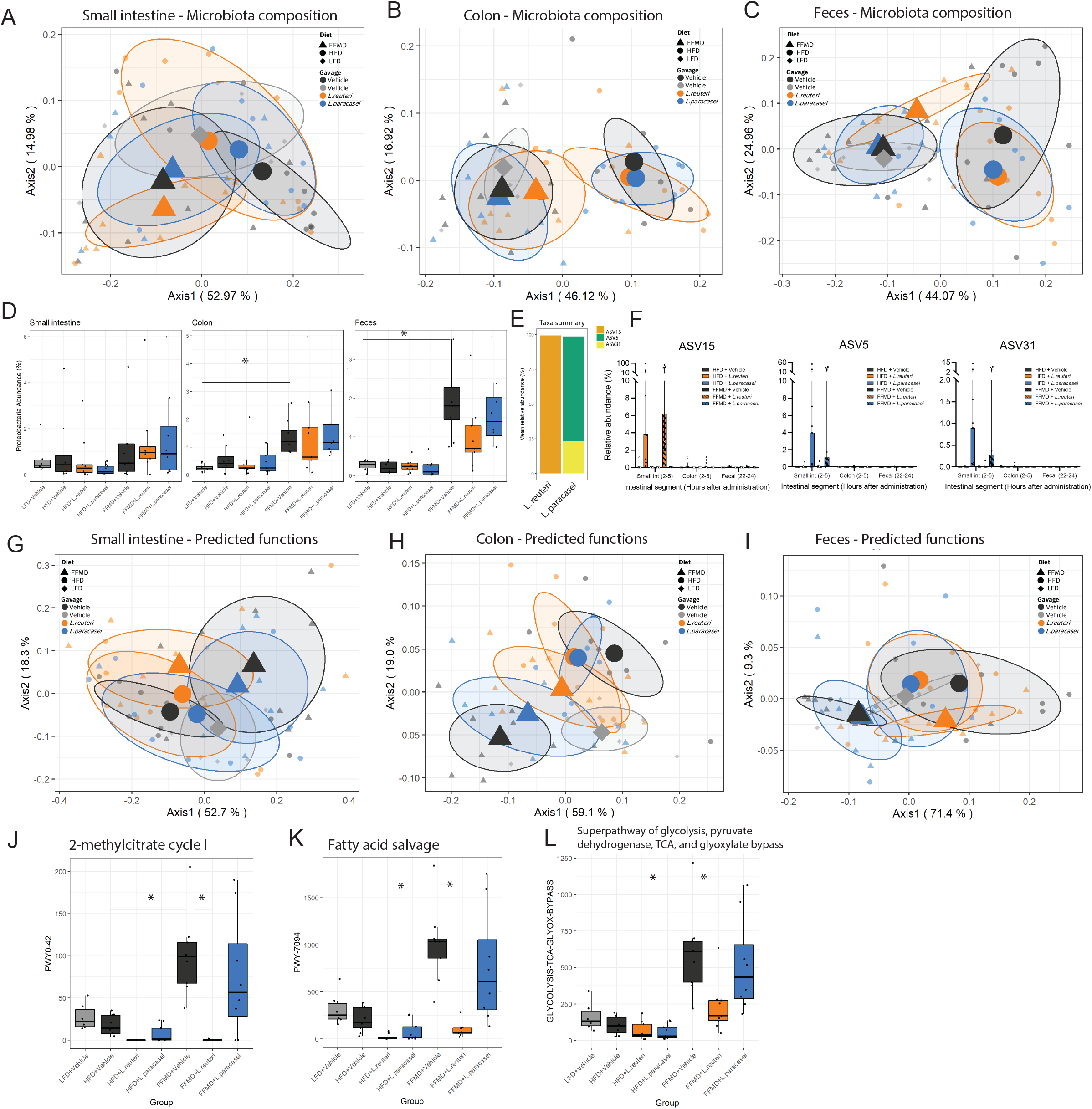
Gut microbiota composition and predicted function are affected by *L. reuteri* in combination with FFMD. **A)** Principal coordinate analysis (PCoA) of microbiota composition using weighted UniFrac distances of small intestine content sampled at the end of Study 1. Centroids indicate group average illustrated with 50% CI and smaller points indicate individual data points. **B)** As a for colon content. **C)** As a for fecal samples collected at week 12 of the study. **D)** Relative abundance in % of *Proteobacteria* phylum in small intestine-, colon content, and fecal samples at the end of Study 1. Points indicate individual data points. Asterisks indicate p-values < 0.05 comparing the indicated groups. **E)** Distribution of amplicon sequence variants (ASVs) as relative abundances identified in *Limosilactobacillus reuteri* DSM 32910 (*L. reuteri*) or *Lacticaseibacillus paracasei* subsp. *paracasei* DSM 32851 (*L. paracasei*) stocks. **F)** Relative abundance in % of ASV15, ASV5 and ASV31, respectively, from the administered *Lactobacillus* strains in small intestine-, colon content, and fecal samples collected 2-5h (small intestine and colon) or 22-24h after latest oral gavage at the end of the study. Points indicate individual data points and bars group mean with interquartile range. **G)** PCoA of predicted functions of the microbiota by KEGG orthology using Bray-Curtis distances of small intestine content sampled at the end of Study 1. Centroids indicate group average illustrated with 50% CI and smaller points indicate individual data points. **H)** As g for colon content. **I)** As g for fecal samples collected at week 12 of the study.-**J)** Relative changes in predicted 2-methylcitrate cycle I BioCyc ID PWY0-42 in fecal samples from the end of Study 1. **K)** As j for fatty acid salvage BioCyc ID PWY-7094. **L)** As j for glycosis pathway BioCyc ID GLYCOLYSIS-TCA-GLYOX-BYPASS. **A-D, F-L)** LFD+Vehicle Small int. and fecal n=6 colon n= 9, HFD+Vehicle Small int. and fecal n=9 colon n=8, HFD+L.reuteri n=9, HFD+L.paracasei n=9, FFMD+Vehicle n=9, FFMD+L.reuteri n=9, FFMD+L. paracasei n=8.

**Table 2:** PERMANOVA p-values

|  | Microbiota composition |  |  | Predicted functions |  |  |
| --- | --- | --- | --- | --- | --- | --- |
|  | Small intestine | Colon | Feces | Small intestine | Colon | Feces |
| HFD+Vehicle vs FFMD+Vehicle | **0.008 \*\*** | **0.001 \*\*\*** | **0.004 \*\*** | **0.006 \*\*** | **0.017 \*** | **0.009 \*\*** |
| HFD+Vehicle vs HFD+*L. reuteri* | 0.20 | 0.82 | 0.30 | 0.47 | 0.84 | 0.26 |
| HFD+Vehicle vs HFD+*L. paracasei* | 0.42 | 0.62 | 0.69 | 0.36 | 0.57 | 0.17 |
| FFMD+Vehicle vs FFMD+*L. reuteri* | 0.14 | 0.08 | 0.10 | **0.04 \*** | **0.01 \*** | **0.04 \*** |
| FFMD+Vehicle vs FFMD+*L. paracasei* | 0.64 | 0.82 | 0.88 | 0.55 | 0.44 | 0.99 |

The tested probiotics did not significantly alter the microbiota composition in small intestine or colon content sampled 2-5h after the latest oral gavage (**Fig 4A-B, Table 2**). The fecal microbiota, sampled 22-24h after the latest gavage, was however transiently modified by *L. paracasei* in HFD-fed mice (**Fig S4A, D**). *L. reuteri* temporarily changed FFMD-induced fecal microbiota composition (**Fig S4B, D**), which tended to be modified by the end of the study (**Fig 4A-C, Table 2)** and was significantly modified in a replicated study (**Fig S4F-H**). Investigation of differential abundances at all phylogenetic levels robustly modified by diet in the different studies revealed an increase in abundance of the phylum *Proteobacteria* by FFMD in colonic and fecal samples compared to LFD (p < 0.001, **Fig 4E, S4I-J**). The FFMD-induced *Proteobacteria* abundance in the distal intestinal segments was numerically reduced by *L. reuteri* and to a lesser degree *L. paracasei* in reproduced studies (**Fig 4E, S4I-J**).

To investigate whether colonization of the administered LAB strains in the intestinal tract was affected by diet we assessed the ASVs originating from the administered *L. reuteri* (ASV15) and *L. paracasei* (ASV5 and ASV31, **Fig 4F**). As expected, these ASVs were found in the small intestine of mice 2-5 hours after receiving the corresponding strain (**Fig 4G**). No change from background signal was detected in colon content 2-5 hours, or feces 22-24 hours after oral gavage (**Fig 4G, Fig S4N**), indicating negligible colonic colonization by the applied probiotics.

We next assessed the predicted functions of the gut microbiota by assessing KEGG orthology. Interestingly, while colon and fecal microbiota composition reflected ingested nutrients (**Fig 4A-C**), the predicted microbiota functionality more closely resembled the pathology of experimental mice (**Fig 4G-I, Table 3**). As such, LFD-fed mice clustered in close vicinity to HFD-fed mice, despite being fed a diet matched to the FFMD (**Table 1, Fig 4G-I**). This observation was further supported by *L. paracasei* treated HFD-fed mice who exhibited an intermediary disease phenotype and likewise clustered between the LFD-fed mice and their vehicle-treated HFD-fed counterparts (**Fig 4G-I**). Despite the lack of clear and reproducible effects on gut microbiota composition from the gavage by the LAB strains there was a significant change in FFMD-induced predicted functions of the microbiota by *L. reuteri* in all assessed intestinal segments at termination (**Fig 4G-I, Table 2**), again mirroring their partial protection against FFMD-induced liver pathology. The specific predicted pathways of fecal samples affected by *L. reuteri* included a reduction in the methylcitrate cycle (**Table 3**, **Fig 4L**) transforming propionic acid to pyruvate and succinate. Functionally, we did not detect differences in cecal propionic acid or other short-chain fatty acids (SCFAs) from the administered strains (**Fig S4O-P**). Interestingly, we observed additional changes in predicted functional pathways in fecal samples by *L. reuteri* including a normalization in fatty acid salvage and glycolysis pathways, which were most pronounced in FFMD-fed mice (**Fig 4K-L, Table 3**).

**Table 3:** Differences in predicted pathways of fecal microbiota week 12 in Study 1 (Top 20)

| HFD+Vehicle vs FFMD+Vehicle |  |  |
| --- | --- | --- |
|  | Pathway | Description |
| 1 | LEU-DEG2-PWY | L-leucine degradation I |
| 2 | PWY0-1479 | tRNA processing |
| 3 | HEMESYN2-PWY | Heme biosynthesis II (anaerobic) |
| 4 | PWY0-1261 | Anhydromuropeptides recycling |
| 5 | P105-PWY | TCA cycle IV (2-oxoglutarate decarboxylase) |
| 6 | PWY-5188 | Tetrapyrrole biosynthesis I (from glutamate) |
| 7 | PWY-5189 | Tetrapyrrole biosynthesis II (from glycine) |
| 8 | GLYCOLYSIS-TCA-GLYOX-BYPASS | Superpathway of glycolysis, pyruvate dehydrogenase, TCA, and glyoxylate bypass |
| 9 | TCA-GLYOX-BYPASS | Superpathway of glyoxylate bypass and TCA |
| 10 | PWY0-42 | 2-methylcitrate cycle I |
| 11 | PWY-5747 | 2-methylcitrate cycle II |
| 12 | PWY-7094 | Fatty acid salvage |
| 13 | PWY-6876 | Isopropanol biosynthesis |
| 14 | PWY-7234 | Inosine-5'-phosphate biosynthesis III |
| 15 | PWY-7328 | Superpathway of UDP-glucose-derived O-antigen building blocks biosynthesis |
| 16 | PWY-7315 | dTDP-N-acetylthomosamine biosynthesis |
| 17 | PWY-621 | Sucrose degradation III (sucrose invertase) |
| 18 | FASYN-INITIAL-PWY | Superpathway of fatty acid biosynthesis initiation (E. coli) |
| 19 | PWY-5971 | Palmitate biosynthesis II (bacteria and plants) |
| 20 | PWY-5989 | Stearate biosynthesis II (bacteria and plants) |
| FFMD+Vehicle vs FFMD+*L.reuteri* |  |  |
|  | **Pathway** | **Description** |
| 1 | LEU-DEG2-PWY | L-leucine degradation I |
| 2 | PWY-5747 | 2-methylcitrate cycle II |
| 3 | PWY-7094 | 2-methylcitrate cycle I |
| 4 | PWY0-42 | 2-methylcitrate cycle I |
| 5 | ALL-CHORISMATE-PWY | Superpathway of chorismate metabolism |
| 6 | DENITRIFICATION-PWY | Nitrate reduction I (denitrification) |
| 7 | PWY-922 | Mevalonate pathway I |
| 8 | PWY-5910 | Superpathway of geranylgeranyldiphosphate biosynthesis I (via mevalonate) |
| 9 | ASPASN-PWY | Superpathway of L-aspartate and L-asparagine biosynthesis |
| 10 | P105-PWY | TCA cycle IV (2-oxoglutarate decarboxylase) |
| 11 | P125-PWY | Superpathway of (R,R)-butanediol biosynthesis |
| 12 | ANAGLYCOLYSIS-PWY | Glycolysis III (from glucose) |
| 13 | CALVIN-PWY | Calvin-Benson-Bassham cycle |
| 14 | COA-PWY | Coenzyme A biosynthesis I |
| 15 | GLCMANNANAUT-PWY | Superpathway of N-acetylglucosamine, N-acetylmannosamine and N-acetylneuraminate degradation |
| 16 | GLYCOLYSIS-TCA-GLYOX-BYPASS | Superpathway of glycolysis, pyruvate dehydrogenase, TCA, and glyoxylate bypass |
| 17 | GLYOXYLATE-BYPASS | Glyoxylate cycle |
| 18 | HSERMETANA-PWY | L-methionine biosynthesis III |
| 19 | NONMEVIPP-PWY | Methylerythritol phosphate pathway I |
| 20 | NONOXIPENT-PWY | Pentose phosphate pathway (non-oxidative branch) |

## Discussion

We used mice fed two distinct but energy-matched high fat and high sucrose diets to test the preventive effects of two LAB strains on obesity and related metabolic disturbances. While both HFD and FFMD induced obesity and compromised glucoregulatory capacity in 12 weeks, FFMD led to enhanced ectopic fat distribution to visceral fat and liver. This induced major phenotypic differences between the HFD and FFMD, notably on body weight gain, energy harvesting, glucose homeostasis, enterohepatic circulation of BAs as well as gut microbiota composition and predicted functionality. Accordingly, the greatest diet-specific impact of the customized FFMD was observed on NAFLD features including hepatocellular morphology, liver lipid, as well as inflammatory profiles and fibrosis. The FFMD-induced liver pathology culminated in significant fibrotic remodeling in >75% of the mice, hence contrasting the results in HFD-fed mice where no fibrotic bridging was observed and only 1 of 9 mice exhibited fibrosis expansion of portal areas. These diet-specific effects can all potentially affect the ability of probiotics to alleviate obesity-related dysmetabolism and related diseases such as NAFLD. This has major physiological significance and clinical relevance given that the choice of experimental diets in preclinical investigation, and the dietary habits of subjects enrolled in clinical trials, are thus likely to influence the efficacy of probiotics. This could lead to the dismissal of promising probiotic candidates for mitigating obesity and related metabolic disorders due to false negative results in preclinical trials, a lack of clinical translatability when advancing to clinical phases where dietary macronutrient composition is not specifically controlled for, or - worst case - lead to the endorsement of probiotics with limited or potentially even unfavorable effects in combination with certain diets not adequately recapitulated in preceeding studies ^20,23^.

In our study, *L. paracasei* reduced obesity development associated with diminished NAFLD exclusively in the context of HFD-feeding without changing energy balance. As a result, 45% of the HFD-mice receiving *L. paracasei* had no histopathological signs of NAFLD, which was not prevented in any other HFD-fed mouse in the study. *L. reuteri* on the other hand improved glucose homeostasis and the more severe NAFLD phenotype only in FFMD-fed mice. This was associated with increased plasma FA 18:2 levels, reported to inversely correlate with human hepatocellular carcinoma (HCC)^47^ and a murine NASH-HCC induction ^48^. *L. reuteri* additionally modified the FFMD-induced BA profile by augmenting the amount of conjugated Bas in cecum. Elevated BA levels could point towards increased energy excretion although such indications were not captured by the bomb calometry measures employed in the current study. Augmented secondary, i.e. dehydroxylated, Bas has also recently been shown to enhance the presence of peripherally induced regulatory T cells ^46,49,50^, hence dampening mucosal immunity; a trait that would be beneficial in the context of diet-induced obesity ^51^. Future studies in either microbiota depleted or germ-free mice are thus warranted to elucidate if and how *L. reuteri* induced BA modifications may affect host immunity.

Although it remains uncertain what caused the diet-dependent effects on probiotic efficacy, possible explanations could relate to direct dietary influence where the administered strains rely on utilization of specific macronutrients. This is exemplified by FFMD-feeding leading to major increases in predicted metabolic functions of the gut microbiota; a traits that was normalized by *L. reuteri* in FFMD-fed mice. It should be noted, though, that while the metabolic benefits of *L. paracasei* associated with reduced weight gain and fat accretion in HFD-fed mice, the improved insulin sensitivity of FFMD-fed animals co-treated with *L. reuteri* was not linked to diminished weight gain, strongly pointing towards dissimilar mechanisms in probiotic efficacy. Importantly, the diet-dependent efficacy of probiotic candidates were highly reproducible across studies and persisted after rigorous statistical testing, cf. our strict approaches adjusting for individual studies, study-batches, potential co-caging and random effects, as well as correcting for potential false discovery rates as outlined in the method section.

We also observed that the diet composition, rather than the amount of dietary fat and high glycemic carbohydrates, was the main driver of gut microbial community structures. This was examplified by a reproducible FFMD-induced augmentation of *Proteobacteria* phylum abundances, which has previously been linked to epithelial damage via disruption in anaerobiosis ^52^ and NASH ^53^. Interestingly, we and others previously observed *Proteobacteria* as the dominant bacterial phylum in extra-intestinal tissues of morbidly obese individuals with T2D compared to normoglycemic weight-matched counterparts ^54,55^. Predicted functions of the microbiota, however, followed disease phenotype rather than diet composition. As such, the tested probiotic strains had negligible impact on microbiota composition, but not its predicted functions, in the sampled intestinal segments and showed minimal signs of colonization of the gastro-intestinal tract, supporting previous reports ^56^. In the study from Zmora *et al*, individual responses to probiotics were suggested to originate from genetic or baseline microbiota differences naturally found in humans ^24^. Both factors were accounted for in our studies, where we show dietary differences alone were sufficient in determining probiotic efficacy. The diet-dependent trait of the strains is warranted to interogate in future studies, just as a combination therapy may alleviate diets dependency. To this end, a combination of the *L. reuteri* and *L. paracasei* strains is currently investigated in an ongoing randomized, double-blinded, placebo-controlled clinical trial in prediabetic subjects (ClinicalTrials.gov identifier NCT04767789).

Differences in probiotic efficacy have been reported for *L. helveticus* R0052*, L. plantarum* WCFS1, and *P. copri ^26–29^* when comparing lean, chow-fed mice to obese, Western diet-fed mice. Our study demonstrates that strain-specific efficacies of probiotics on body composition, insulin resistance, and NAFLD were reproducible, yet highly influenced by the type of obesogenic diet, and thus substantially expanding the understanding of dietary influence on probiotic efficacy. While future studies are warranted to elucidate the generic implication of our findings, the dietary interactions described here may likely extend beyond the tested probiotic strains. Potentially, this could aid in explaining empiric observations of responders versus non-responders to probiotics in general and might indicate a need for pre-treatment stratification of patients.

## Limitations

The study was limited by the use of 16S rRNA gene amplicons for evaluation of the gut microbiota with lower resolution of microbial species compared to shotgun sequencing and relying on assumptions to predict microbiota functionality. While the macronutrients of the diets are relatively similar between the HFD and FFMD, the nutrient sources are vastly different. Future lines of research should therefore elucidate the precise dietary components driving the reported phenotypic discrepanices.

## Acknowledgment

We thank Joanie Dupont-Morissette, Christine Dallaire, Jacinthe Julien, Laurence Morin, and Jenny Rancourt-Mercier from IUCPQ, Laval University, for excellent technical assistance for *in vivo* protocols. Additionally, we thank Marie-Julie Dubois and Michael Schwab from IUCPQ for data generation during the manuscript revision process. Sincere gratitude to Lea B. S. Hansen for essential bioinformatic input. Additionally, we thank Torsten Schröder from the Institute of Nutritional Medicine for data generation as well as Jeffrey Schultchen from NZ for providing the LAB strains.

## Data availability statement

The data that support the findings of this study are available in Mendeley Data at

DOI: http://doi.org/10.17632/72j8tkm36g.1

---

**Supplementary Table 1:** Grading system for murine NAFLD activity score, adapted from Liang *et al* ^32^.

| Histological feature | Score |  |  |  |
| --- | --- | --- | --- | --- |
|  | 0 | 1 | 2 | 3 |
| **Steatosis score** |  |  |  |  |
| Macrovesicular steatosis | < 5% | 5-33% | 33-66% | >66% |
| Microvesicular steatosis | < 5% | 5-33% | 33-66% | >66% |
| Hypertrophy | < 5% | 5-33% | 33-66% | >66% |
| **Inflammation Score** |  |  |  |  |
| Number of inflammatory foci per field | <0.5 | 0.5-1.0 | 1.0-2.0 | >2.0 |

**Supplementary Table 2:** Extended staging system for hepatic fibrosis as introduced by Ishak et al. ^33^

| Histological changes | Score |
| --- | --- |
| No fibrosis | 0 |
| Fibrous expansion of some portal areas with or without short fibrous septa | 1 |
| Fibrous expansion of most portal areas with or without short fibrous septa | 2 |
| Fibrous expansion of most portal areas with occasional portal to portal (P-P) bridging | 3 |
| Fibrous expansion of portal areas with marked bridging (P-P as well as portal to central (P-C)) | 4 |
| Marked bridging (P-P and/or P-C) with occasional nodules (incomplete cirrhosis) | 5 |
| Cirrhosis, probable or definite | 6 |

**Supplementary Table 3:**
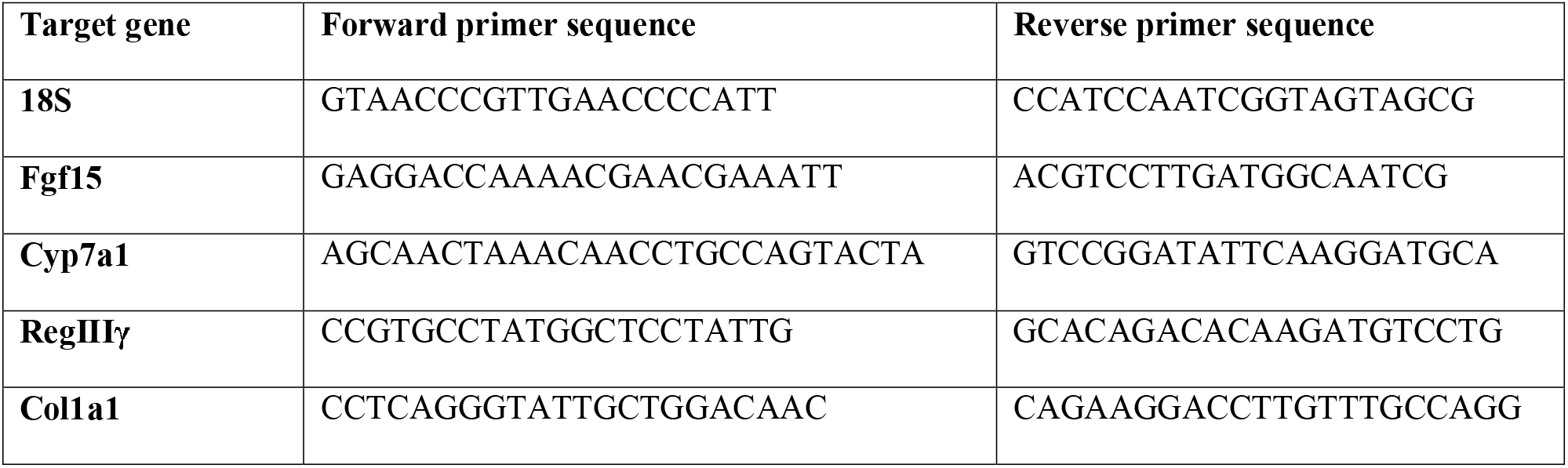
RT-qPCR primer sequences.

**Supplementary figure S1:**
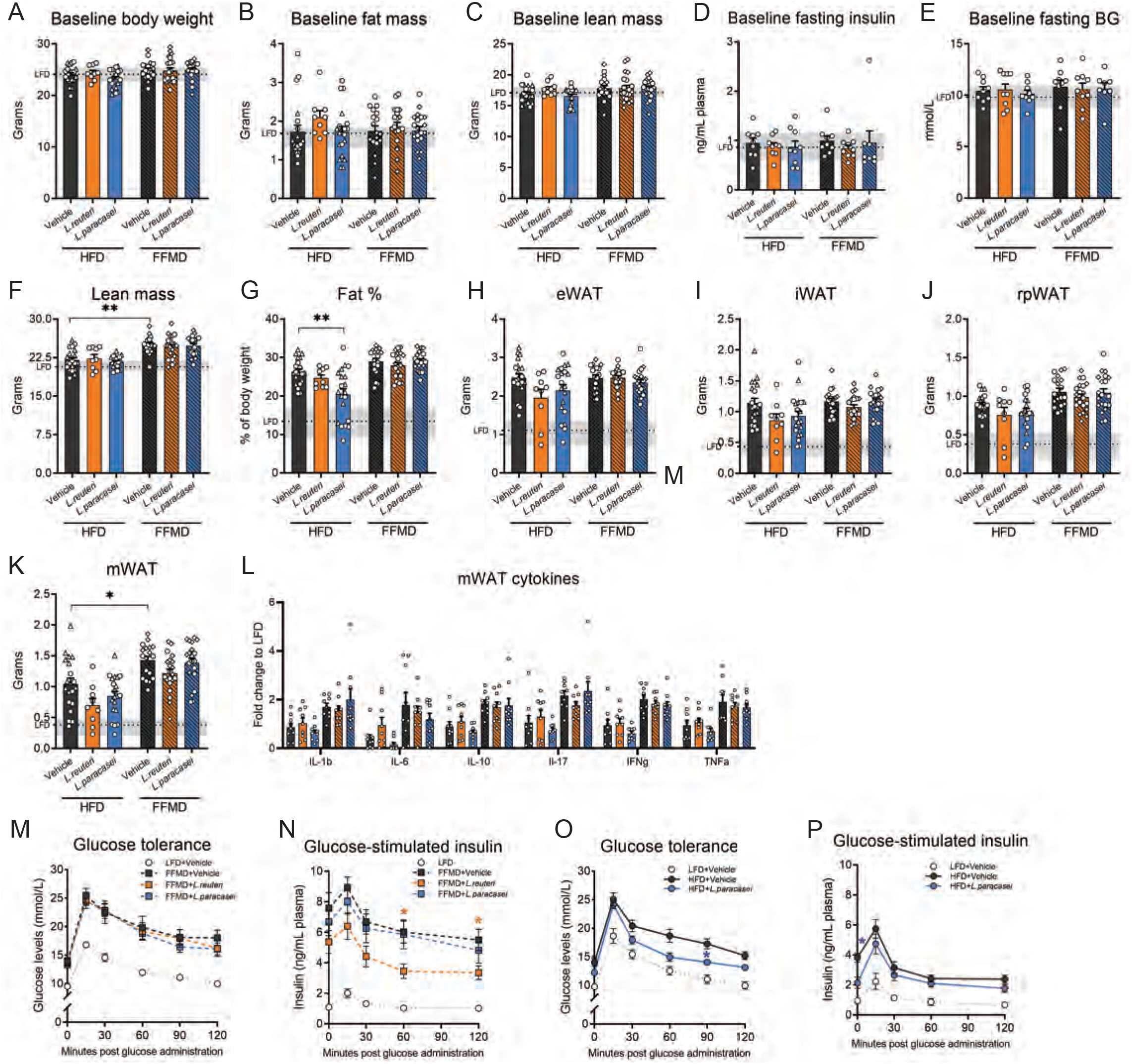
Baseline measures, final adipose tissue weights and replicated glucose tolerance tests. **A)** Baseline body weight in grams. **B)** Baseline fat mass in grams assessed by magnetic resonance (MR) scans. **C)** Baseline lean mass in grams assessed by MR scans. **D)** 5h fasting plasma insulin levels at baseline from Study 1. **E)** 5h fasting blood glucose (BG) levels at baseline from Study 1. **F)** Lean mass in grams assessed by MR scans at week 12 of the study. **G)** Fat percentage assessed by MR scans to total body weight at week 12 of the study. **H)** Weight of epididymal white adipose tissue (eWAT) at the end of the study. **I)** Weight of subcutaneous inguinal WAT (iWAT) at the end of the study. **J)** Weight of visceral retroperitoneal WAT (rpWAT) at the end of the study. **K)** Weight of visceral mesenteric WAT (mWAT) at the end of the study. **L)** Cytokine levels in mWAT tissue normalized to total tissue weight. **M)** Blood glucose values from Study 3 during oral glucose tolerance test (oGTT) with 2 μg/g lean mass glucose in week 10. **N)** Corresponding plasma insulin levels during oGTT shown in l. **O)** Blood glucose values from Study 2 during oral glucose tolerance test (oGTT) with 1.5 μg/g lean mass glucose in week 10. **P)** Corresponding plasma insulin levels during oGTT shown in n. **A-E)** Baseline measures prior study start while all groups were fed LFD and later divided into the indicated groups. **A-P)** Bars or lines indicate group mean ± SEM. Dashed line indicate mean of LFD group with interquartile range shown in grey. **A-K)** Points represent individual data points. Asterisks indicate fdr-corrected q-values <0.05 by linear mixed effects models comparing indicated groups. **A-C, F-K)** Point shape indicates three individual studies where rounds indicate Study 1, triangles Study 2, squares Study 3. LFD+Vehicle n=21, HFD+Vehicle n=20, HFD+L.reuteri n=9, HFD+L.paracasei n=20, FFMD+Vehicle n=18, FFMD+L.reuteri n=21, FFMD+L. paracasei n=20. **D-E)** LFD+Vehicle n=9, HFD+Vehicle n=9, HFD+L.reuteri n=9, HFD+L.paracasei n=9, FFMD+Vehicle n=9, FFMD+L.reuteri n=9, FFMD+L. paracasei n=8. **M-O)** Asterisks indicate p-values <0.05 using two-way ANOVA with multiple comparisons between vehicle-treated groups or LAB group to vehicle-treated group fed the same diet with Bonferroni post-hoc test. **M-N)** LFD+Vehicle n=12, FFMD+Vehicle n=9, FFMD+L. reuteri n=9, FFMD+L. paracasei n=12. **O-P)** LFD+Vehicle n=10, HFD+Vehicle n=10, HFD+L.paracasei n=11.

**Supplementary figure S2:**
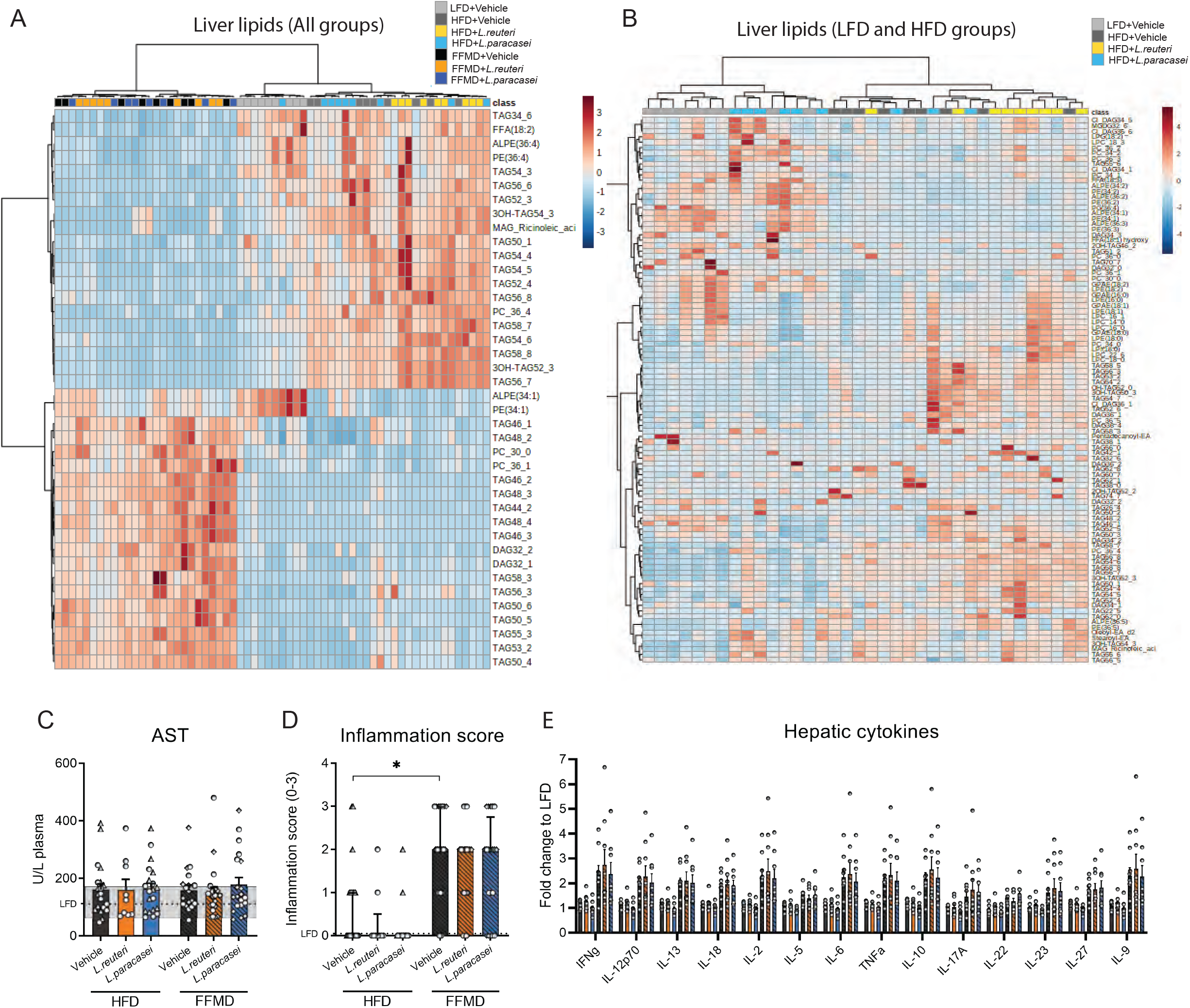
Liver lipids and cytokine levels. **A)** Heatmap of 40 most discriminating lipids in liver tissue from end of study 1 assesed by lipidomics. Experimental group of each sample are indicated by color and distributed by hierarchical clustering. **B)** Heatmap of 100 most discriminating lipids of LFD and HFD fed groups in liver tissue from end of study 1 assesed by lipidomics. Experimental groups are indicated by color distributed by hierarchical clustering. **C)** Aspartate transaminase (AST) levels in plasma at the end of the study. **D)** Histological inflammation score from 0-3 blindly assessed from H&E-stained liver tissue. Asterisks indicate p-values <0.05 by Kruskal-Wallis test with multiple comparisons between vehicle-treated groups or LAB group to vehicle-treated group fed the same diet with Dunn’s post-hoc test.. **D)** Cytokine levels as fold change to LFD reference measured in liver tissue in Study 1. Bars indicate group mean ± SEM with points illustrating individual data points. **C-D)** Bars indicate group mean ± SEM with points illustrating individual data points. Dashed line indicate mean of LFD group with interquartile range shown in grey. Point shape indicates three individual studies where rounds indicate Study 1, triangles Study 2, squares Study 3. **A, E)** LFD+Vehicle n=9, HFD+Vehicle n=9, HFD+L.reuteri n=9, HFD+L.paracasei n=9, FFMD+Vehicle n=9, FFMD+L.reuteri n=9, FFMD+L. paracasei n=8. **B)** LFD+Vehicle n=9, HFD+Vehicle n=9, HFD+L.reuteri n=9, HFD+L.paracasei n=9. **C, D)** LFD+Vehicle n=21, HFD+Vehicle n=20, HFD+L.reuteri n=9, HFD+L.paracasei n=20 (Except for D where n = 18), FFMD+Vehicle n=18, FFMD+L.reuteri n=21, FFMD+L. paracasei n=20.

**Supplementary figure S3:**
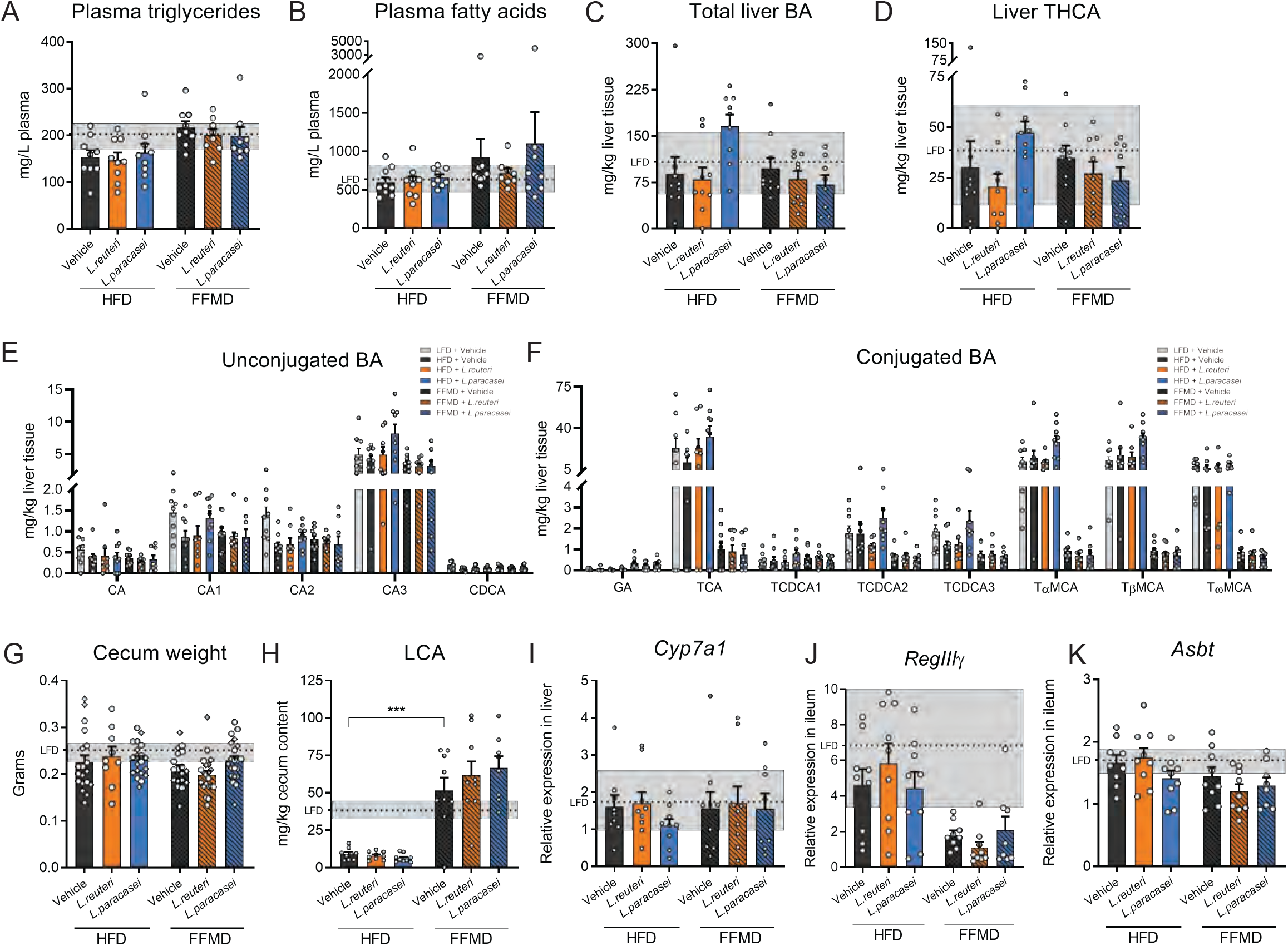
Plasma lipids, liver and cecum bile acids (BA) **A)** Plasma triacylglycerol levels from the end of Study 1. **B)** Plasma free fatty acids (FFA) levels from the end of Study 1. **C)** Total measured BA concentration in liver tissue. **D)** Concentration of 3α,7α,12α-trihydroxycholestanoic acid (THCA) in liver tissue. **E)** Concentration of unconjugated BA, cholic acid (CA) -derivatives and chenodeoxycholic acid (CDCA) in liver tissue. **F)** Concentration of conjugated BA in liver tissue. **G)** Weight of full cecum in grams. Points represent individual mice in three individual studies indicated by point shape where rounds indicate Study 1, triangles Study 2, squares Study 3. LFD+Vehicle n=21, HFD+Vehicle n=19, HFD+L.reuteri n=9, HFD+L.paracasei n=20, FFMD+Vehicle n=18, FFMD+L.reuteri n=21, FFMD+L. paracasei n=20. **H)** Concentration of secondary bile acid lithocholic acid (LCA) in cecum content. **I)** Relative cholesterol 7 alpha-hydroxylase (Cyp7a1) gene expression in liver tissue by RT-qPCR. **J)** Relative apical sodium-bile acid transporter (Asbt) gene expression in ileum tissue by RT-qPCR. **K)** Relative Regenerating islet-derived protein 3-gamma (RegIIIγ) gene expression in ileum tissue by RT-qPCR. **A-K)** Bars indicate group mean ± SEM. Asterisks indicate fdr-corrected q-values <0.05 using linear mixed effects models comparing indicated groups. **A-D, G-K)** Dashed line indicates mean of LFD group with interquartile range shown in grey. **A-B)** LFD+Vehicle n=8, HFD+Vehicle n=9, HFD+L.reuteri n=9, HFD+L.paracasei n=9, FFMD+Vehicle n=9, FFMD+L.reuteri n=8, FFMD+L. paracasei n=8. **C-D)** LFD+Vehicle n=9, HFD+Vehicle n=9, HFD+L.reuteri n=9, HFD+L.paracasei n=9, FFMD+Vehicle n=9, FFMD+L.reuteri n=9, FFMD+L. paracasei n=8. **E-F, H-K)** LFD+Vehicle n=8, HFD+Vehicle n=9, HFD+L.reuteri n=8, HFD+L.paracasei n=9, FFMD+Vehicle n=9, FFMD+L.reuteri n=8, FFMD+L. paracasei n=8.

**Supplementary figure S4:**
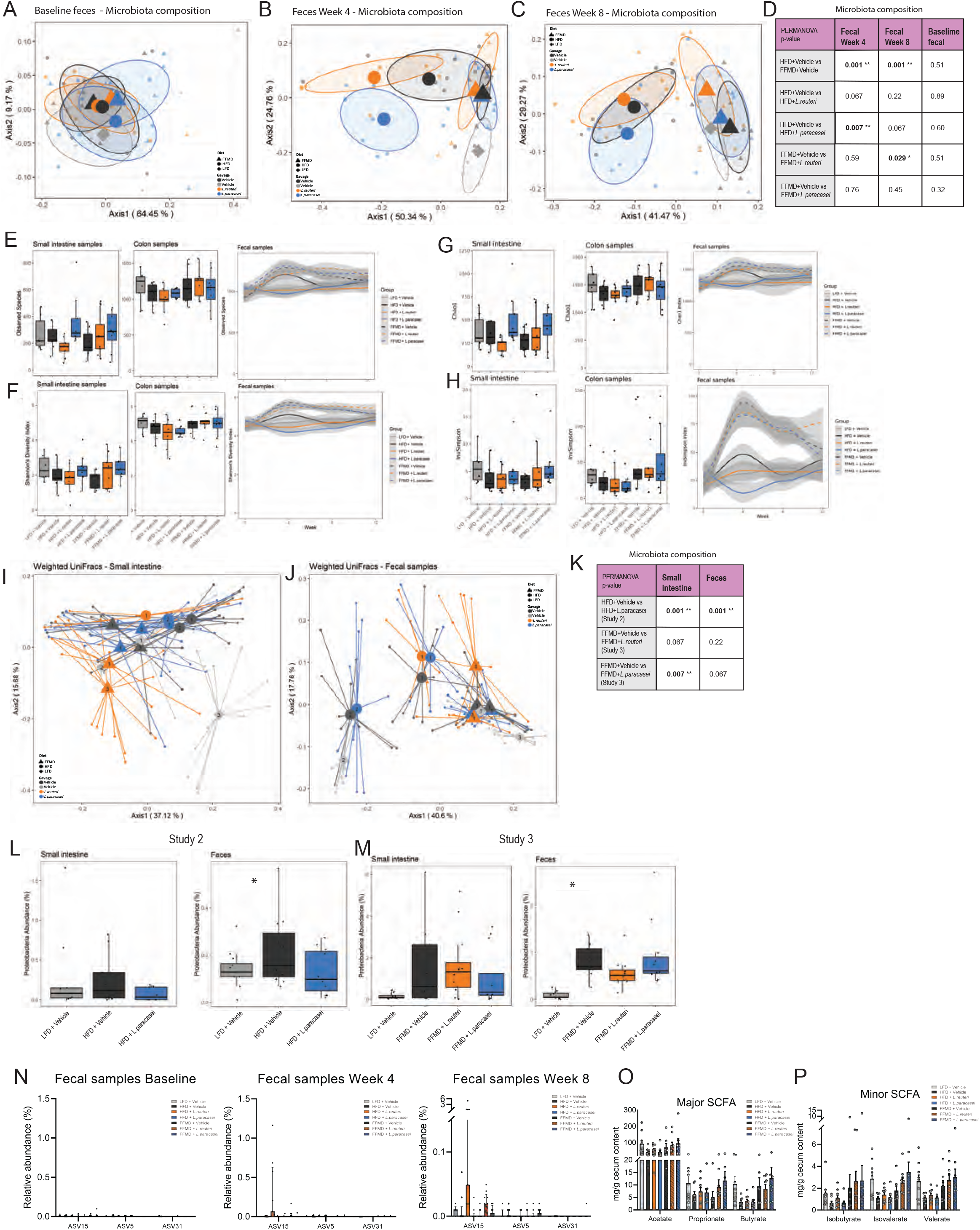
Baseline microbiota composition and alpha diversity assessments. **A)** Principal coordinate analysis (PCoA) of baseline fecal samples collected prior to the start of Study 1 when all mice were fed LFD. Centroids indicate group average illustrated with 50% CI and smaller points indicating individual data points of the later designated experimental groups. **B) PCoA** of microbiota composition using weighted UniFrac distances of fecal samples collected in week 4 of Study 1. Centroids indicate group average illustrated with 50% CI and smaller points indicating individual data points. **C)** As b for fecal samples collected week 8. **D)** PERMANOVA p-values comparing the microbiota composition between indicated groups of Study 1 on weighted UniFrac distances. Asterisks indicate p-values < 0.05. **E)** Alpha diversity measure of observed species richness in small intestine and colon content from the end of the study as well as in fecal samples over time. **F)** As e with Shannon’s Diversity index. **G)** As e with Chao1 index. **H)** As e using Inverse Simpson index. **I)** PCoA of small intestine samples in all three studies indicated by number in the centroid. Centroid indicates group average per study projecting to individual data points. **J)** As i for fecal samples collected at the end of the three studies. **K)** PERMANOVA p-values comparing the microbiota composition between indicated groups in individual studies on weighted UniFrac distances. Asterisk indicate p-values < 0.05. **L)** Relative abundance in % of phylum *Proteobacteria* in fecal samples and small intestine content at the end of Study 2. Boxplot indicates distribution within the groups and points indicate individual data points. Asterisk indicate p-value < 0.05 comparing the indicated groups. LFD+Vehicle Small int n=9 and feces n=12, HFD+Vehicle Small int n=9 and feces n=11, HFD+L. paracasei Small int n=9 and feces n=11. **M)** As i for Study 3. LFD+Vehicle n=12, FFMD+Vehicle n=9, FFMD+L. reuteri n=12, FFMD + L. paracasei Small int n=12 and feces n=11. **N)** Relative abundance in % of ASV15, ASV5 and ASV31 from the administered LAB strains in fecal samples collected respectively at baseline prior any administration of the strains, in week 4 and week 8 of Study 1. Week 4 and 8 sampling was carried out 22-24h after latest oral. Points indicate individual data points and bars group mean with interquartile range. **O)** Major short-chain fatty acids (SCFA). **P)** Minor short-chain fatty acids SCFA. **A-D, N)** LFD+Vehicle n=5 (except for baseline where n=6), HFD+Vehicle n=9, HFD+L.reuteri n=9, HFD+L.paracasei n=9, FFMD+Vehicle n=9, FFMD+L.reuteri n=9, FFMD+L. paracasei n=8. **E-H)** LFD+Vehicle Small int. and fecal n=6 colon n= 9, HFD+Vehicle Small int. and fecal n=9 colon n=8, HFD+L.reuteri n=9, HFD+L.paracasei n=9, FFMD+Vehicle n=9, FFMD+L.reuteri n=9, FFMD+L. paracasei n=8. **I-J)** LFD+Vehicle n = 18, HFD+Vehicle n=18, HFD+L.reuteri n=9, HFD+L.paracasei Small int n=18 and feces n=20, FFMD+Vehicle n=18, FFMD+L.reuteri n=21, FFMD+L. paracasei Small int n=20 and feces n=19. **O-P)** LFD+Vehicle n=9, HFD+Vehicle n=8, HFD+L.reuteri n=9, HFD+L.paracasei n=9, FFMD+Vehicle n=7, FFMD+L.reuteri n=9, FFMD+L. paracasei n=8.

